# Sialidase variability in *Gardnerella*: genetics, taxonomy, function, and clinical presentation

**DOI:** 10.64898/2026.01.10.698787

**Authors:** Marie Vasse, Nicola Mayrhofer, Valerie Masete, Jo-Ann S. Passmore, Hélène Jean-Pierre, Sylvain Godreuil, Rémy Froissart, Ignacio G. Bravo

## Abstract

Bacterial vaginosis (BV) is the most common vaginal condition among women of reproductive age. Despite over six decades of research, the pathogenesis and etiology of BV remain largely unknown. Members of the *Gardnerella* genus are common components of the healthy vaginal microbiota, although their uncontrolled proliferation plays a key role in BV pathogenesis. Recent taxonomic revisions have highlighted extensive genetic and functional diversity among *Gardnerella* spp., but the relationship between genetics, taxonomy, and virulence remains elusive. Sialidases are key virulence factors in BV, degrading the protective epithelial mucus layer, facilitating colonization and releasing carbon and nitrogen sources. However, robust clinical evidence linking sialidases to increased BV risk remains limited. Here, we analyzed clinical *Gardnerella* isolates from BV-positive (75%) and BV-negative (25%) samples (n = 66) collected at the University Hospital of Montpellier, France. We inferred phylogenetic relationships, characterised the *nanH* sialidase gene repertoire, quantified *in vitro* sialidase activity, and evaluated antibiotic resistance profiles. Most isolates belonged to *Gardnerella* clades 1, 2, and 4 (63.6%, 19.7%, and 16.7%, respectively). The three *nanH* genes were unevenly distributed across clades: *nanH1* was most common (∼80%), highly prevalent in clades 1-3; *nanH2* was rare (6%); and *nanH3* (∼38%) was predominantly found in clade 2 - typically co-occurring with *nanH1*. Sialidase activity varied both between and within clades, and was only weakly associated with the *nanH* gene repertoire. Neither taxonomy nor sialidase-associated traits were reliable predictors of BV status, and we found no clear link between taxonomy and antibiotic resistance. Together, these results reveal substantial discordance in the genotype-phenotype relationship in *Gardnerella*, suggesting epigenetic, physiological and/or ecological modulation. Our findings highlight the need for a better understanding of bacterial physiology, plasticity, and ecology to explain microbial traits of clinical importance.

## Introduction

Bacterial vaginosis (BV) is the most common vaginal condition among women of reproductive age, with prevalence rates ranging from 23% to 29% across global regions [1]. BV is characterized by the replacement of bacteria that produce lactic acid in the healthy vaginal microbiota (primarily Lactobacilli) by a polymicrobial consortium of facultative and strict anaerobes (including *Gardnerella vaginalis* and *Prevotella bivia*) [2–6]. Changes in microbiota composition are associated with several health issues such as an increased incidence of sexually transmitted infections and an increased risk of preterm birth or infertility [7–9]. Despite more than six decades of research, the natural history of BV remains incompletely characterized. The etiological perspective on BV has shifted from an initial mono-microbial view with *G. vaginalis* as the sole culprit [10], towards the current ecological perspective as a dysbiosis of the vaginal microbiota [11]. The detection of *G. vaginalis* in vaginal samples from both BV-positive and BV-negative women [2, 12, 13] has generated interest in characterizing the diversity among *Gardnerella* isolates and in determining whether genotypic and/or phenotypic variation underlies the differing associations between specific vaginal microbiota profiles and clinical presentation.

Bacterial sialidase activity has been proposed as a key driver in the development of BV [14] and is therefore a strong candidate marker of BV-associated dysbiosis. Sialidases are enzymes that cleave terminal sialic acid molecules from glycosylated substrates. Sialylated structures are very common on the vaginal surface and secretions, with a proposed protective and immunological role [15]. By liberating host-derived nutrients, bacterial sialidases can be regarded as communal resources, benefiting co-resident microbes that lack the enzyme themselves. Elevated levels of sialidase activity in vaginal fluids of women with BV were first reported three decades ago [16], and subsequent studies have confirmed the association between sialidase activity and BV [17–19], supporting the evaluation of vaginal sialidase levels as a biomarker for BV diagnosis [19–22]. In 2013, Lewis *et al.* showed that cultured isolates of *G. vaginalis* hydrolyze and deplete mucus components containing sialic acid in the vagina and characterized the sialic acid catabolic gene cluster [23]. In vaginal samples, the high prevalence of the *G. vaginalis* putative sialidase gene, initially called *sialidase A* and now known as *nanH1*, has later been associated with BV and biofilm formation in clinical samples [24]. Not all *Gardnerella* isolates, however, encode *nanH1* [24–26] and not all isolates carrying the gene exhibit sialidase activity [25, 27–29]. More recently, Robinson and collaborators reported little or no enzyme activity of the *nanH1* gene product against most tested substrates, while two newly described sialidase genes, *nanH2* and *nanH3*, effectively released terminal sialic acids from nearly all substrates tested [28]. The authors thus concluded that *nanH2* and *nanH3* were the main sources of sialidase activity produced by *Gardnerella* spp. isolates. Subsequent studies have largely supported this hypothesis, highlighting some inconsistencies in the association between the presence of *nanH2* or *nanH3* and measured sialidase activity [29–31].

Early evaluations of *G. vaginalis* diversity were based on enzyme activity biotyping methods [32, 33] as well as genotype differentiation [34], and shed light on a wide range of both phenotypes and genotypes. In the following years, many culture-based studies further highlighted a broad range of variable phenotypic traits in *G. vaginalis* [27, 35–39]. In a more recent attempt to characterize the genotypic diversity of *G. vaginalis*, Jayaprakash and colleagues [40] defined four subgroups (A to D) based on differences in the universal target region of the chaperonin-60 (*cpn60* ) gene. In the same year, Ahmed *et al.* proposed a classification of *G. vaginalis* into four clades (1 to 4) based on whole-genome compositional data [41]. These four clades harbored distinct gene pools and genomic properties and were associated with three different ecotypes by Cornejo *et al.* [42]. Both classifications [40, 41] are largely congruent and were later reconciled by Schellenberg *et al.* [25]. Finally, in 2019, in a study based on 81 whole-genome sequences of *Gardnerella* isolates, Vaneechoutte and colleagues proposed to split the genus into four newly defined species (namely *Gardnerella vaginalis*, *Gardnerella piotii*, *Gardnerella leopoldii*, and *Gardnerella swidsinskii*) and nine additional ”genome species” that were not named nor formally described [43]. Two of these ’genome species’ (3 and 8) were later named as *Gardnerella pickettii* and *Gardnerella greenwoodii*, respectively [44]. Table1 summarizes the relationship between the different classifications.

Castro and colleagues highlighted the lack of strict correspondence between taxonomic stratification among *Gardnerella* species and BV clinical presentation [45]. Globally, clades 1 and 2 are typically associated with BV, while the association with BV for clades 3 and 4 would be less common [12, 40, 46–49]. In addition, multiple *Gardnerella* clades/species often coexist in the same vaginal microbiota [12, 40, 46–50], and BV-positive samples are associated with a higher likelihood of detecting multiple *Gardnerella* clades compared to BV-negative samples [12, 46, 47, 49, 50]. Many of these studies, however, report only the presence or absence of individual *Gardnerella* clades and do not provide information on relative abundances. In contrast, Hill and colleagues investigated the relative abundances of each species in vaginal samples and showed that, although co-occurrence was common, most microbiota were strongly dominated by a single *Gardnerella* species [51].

Experimental evaluation and quantification of the sialidase activity produced by *Gardnerella* spp. shows a wide range of variation between isolates *in vitro* [16, 23, 25–29, 31, 36, 43] (See Summary Table S9). A few studies have investigated the prevalence of sialidase activity in different *Gardnerella* clades and generally report that all isolates from clade 2 (with *G. piotii* and *G. pickettii* ) and some isolates from clade 1 (with *G. vaginalis* and genome species 2) exhibit sialidase activity. In contrast, isolates from clades 3 (with *G. greenwoodii* and genome species 9 and 10) and 4 (with *G. leopoldii* and *G. swidsinskii*) show little or no sialidase activity [25, 29, 31, 43]. This taxonomic pattern of sialidase activity represents an important step towards elucidating the link between *Gardnerella* taxonomy and the observed diversity in sialidase activity between isolates. However, studies examining the taxonomic distribution of sialidase activity have largely lacked clinical context with respect to BV [25, 29, 31, 43, 44], whereas studies linking sialidase activity with BV status typically do not resolve *Gardnerella* taxonomy ([16, 23, 36]). In a recent synthesis of *Gardnerella* taxonomic diversity, Castro et al. further highlighted the outstanding question of whether isolates from women with clinically diagnosed BV differ from those in BV-negative women in their susceptibility to antimicrobial agents [45].

To our knowledge, no study has simultaneously linked *Gardnerella* taxonomy, *nanH* gene repertoire, antibiotic susceptibility, and quantitative sialidase activity in clinically characterized BV samples. In the present work, we address this important gap by integrating *Gardnerella* spp. taxonomy - based on housekeeping and *nanH* genes - with measurements of functional traits in the context of clinical BV status. Using 66 clinical *Gardnerella* isolates from BV-negative and BV-positive samples, we first characterized the repertoire of *nanH* sialidase genes and determined susceptibility to four commonly used antibiotics. We then assessed phylogenetic relationships based on the *nanH1* gene and the *cpn60* marker. Finally, for a subset of 40 isolates (10 BV-negative and 30 BV-positive), we quantified secreted sialidase activity from colonies grown on solid medium.

## Results

### Phylogenetic analysis identifies five *Gardnerella* clades, with clinical isolates enriched in clades 1, 2, and 4

Following isolation of 66 clinical strains and retrieval of one representative genome per *Gardnerella* species, we examined their phylogenetic relationships using the conserved *cpn60* marker. Our maximum likelihood phylogenetic analysis confirms five well-defined *Gardnerella* lineages (Figure 1). Concordance with previous taxonomic classifications is shown in Table 1. We resolved the taxonomic placement of several different *Gardnerella* genome species previously designated *incerta sedis*. *Gardnerella* genome species 2 clustered as the sister taxon of *G. vaginalis* reference strains from both the American (ATCC 14018) and German (DSM 4944) type culture collections, with all three belonging to clade 1. *Gardnerella* genome species 11 grouped as the sister taxon of *G. piotii* and *G. pickettii*, together forming clade 2. *G. greenwoodii* and *Gardnerella* genome species 9 and 10 were closely related and clustered within clade 3. *Gardnerella* genome species 13 was identified as the sister taxon of *G. leopoldii*, *G. swidsinskii*, and of *Gardnerella* genome species 7, with all four belonging to clade 4. Finally, *Gardnerella* genome species 12 did not cluster with any other isolate and constituted the sole representative of clade 5 (Figure 1). Notably, although *Gardnerella* genome species 7 clusters confidently within clade 4, it exhibits an unusually large branch distance from other members of the clade based on the *cpn60* marker (Supplementary Figure S3).

**Figure 1:**
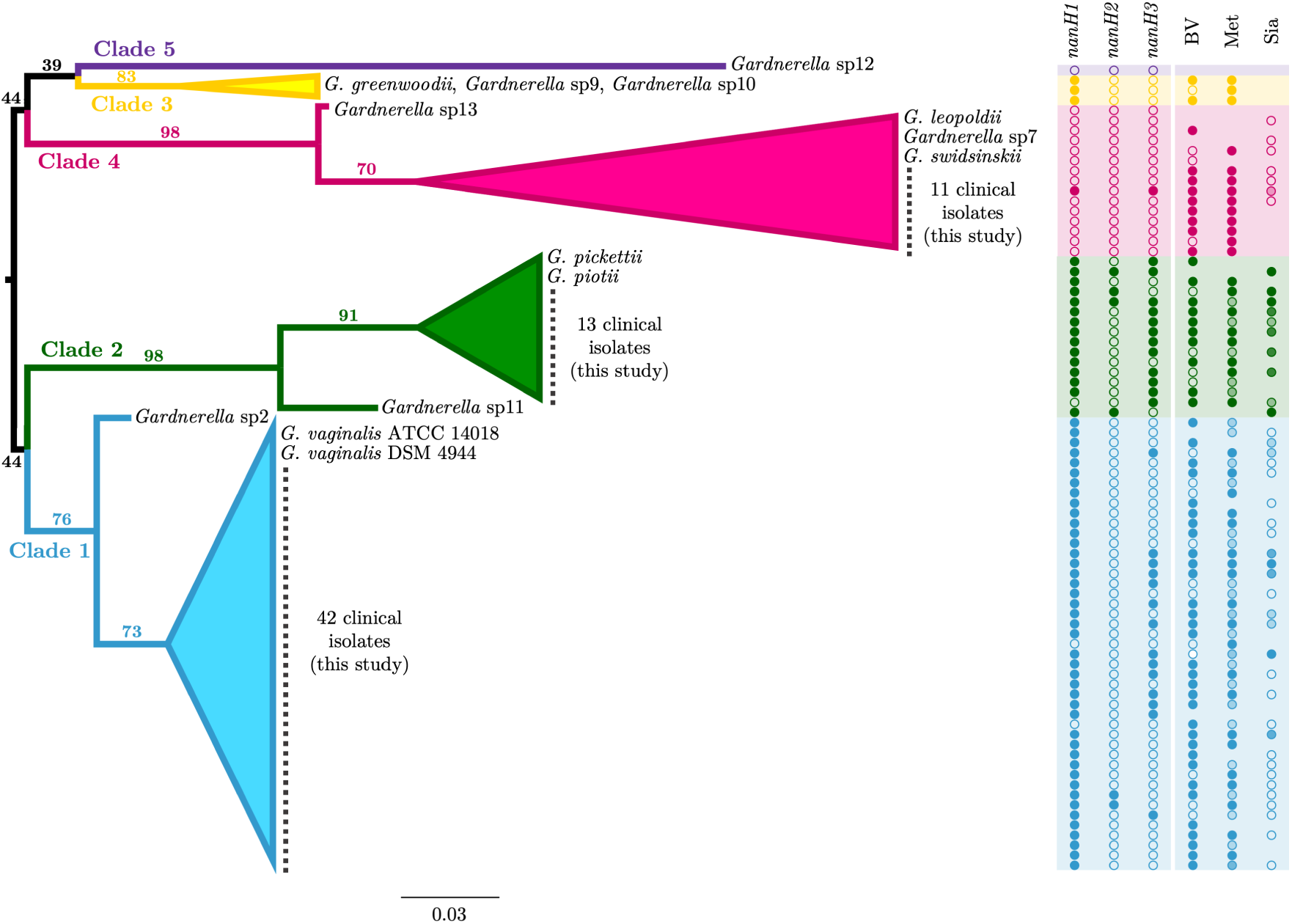
**Phylogenetic reconstruction and associated genotypic and phenotypic features of *Gardnerella* spp. isolates**. (left) Mid-point rooted, maximum-likelihood best-known phylogenetic tree based on *cpn60* gene sequences. Branch support values are indicated and derived from 1,000 bootstrap replicates. Scale for branch length corresponds to number of nucleotide substitutions per site. Colored labels represent the five major *Gardnerella* lineages proposed for taxonomic classification. Clinical isolates from this study are marked as small black squares within the corresponding clades. (right) Overview of genotypic and phenotypic features associated with each isolate. *nanH1-3* : Presence (filled circle) or absence (empty circle) of the three sialidase genes, determined by PCR amplification (for study isolates) or BLAST (for published genomes). *BV* : Bacterial vaginosis status (positive = filled circle; negative = empty circle), based on Spiegel’s criteria applied to the vaginal sample microbiota. *Met* : Resistance to metronidazole, assessed by E-test. Circle transparency reflects MIC values ranging from 1 to 256 µg/mL. *Sia*: Sialidase activity levels measured by fluorometric assay. Transparency reflects activity levels (high, medium, low, or none). For published genomes, BV status was inferred from Nugent scores (positive: 7–10, negative: 0–6) when available, or based on author classifications. *Sia* data was treated as a binary variable: positive (filled circle) or negative (open circle). *Met* was extracted from corresponding publications and visualized using the same transparency scale as for clinical isolates. Literature sources for data in *BV*, *Met*, and *Sia* columns are provided in Supplementary Table S2.

**Table 1:**
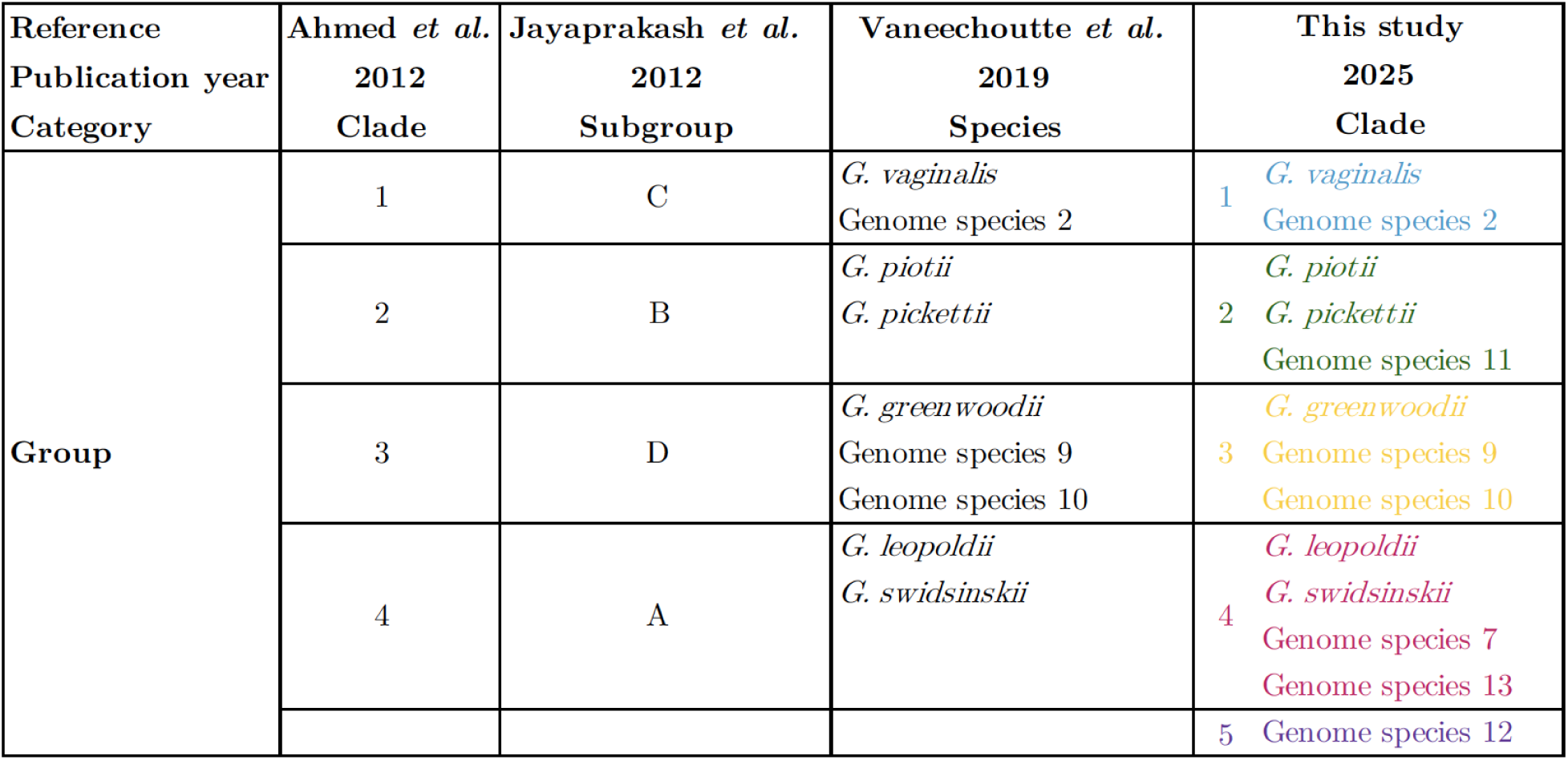
Taxonomy within the *Gardnerella* complex. . The table illustrates the alignment between the previously described four clades and subgroups of *G. vaginalis* (first two columns) and the newly described *Gardnerella* species (last two columns). Clade numbering (1–5) and associated colour coding are used consistently throughout the text.

Based on *cpn60* sequences, the majority of our clinical isolates (42 out of 66) were assigned to clade 1, followed by 13 isolates in clade 2, and 11 in clade 4. This distribution of isolates across phylogenetic clades is broadly consistent with three previous studies [29, 31, 43], with no significant differences in isolate distribution after correction for multiple comparisons (multinomial tests p-values: p = 0.099, p = 0.057 and p = 0.057, respectively) (Figure 2). Other studies reported differing patterns (multinomial tests: all p *<* 0.001). Schellenberg *et al.* 2016 [25] observed a more balanced distribution among clades 1, 2, and 4 while Jayaprakash *et al.* [40] and Ahmed *et al.* [41] reported a predominance of clade 2 and 3, respectively. Vaneechoutte *et al.* [43] identified several genomes that did not cluster within the four previously described clades.

**Figure 2:**
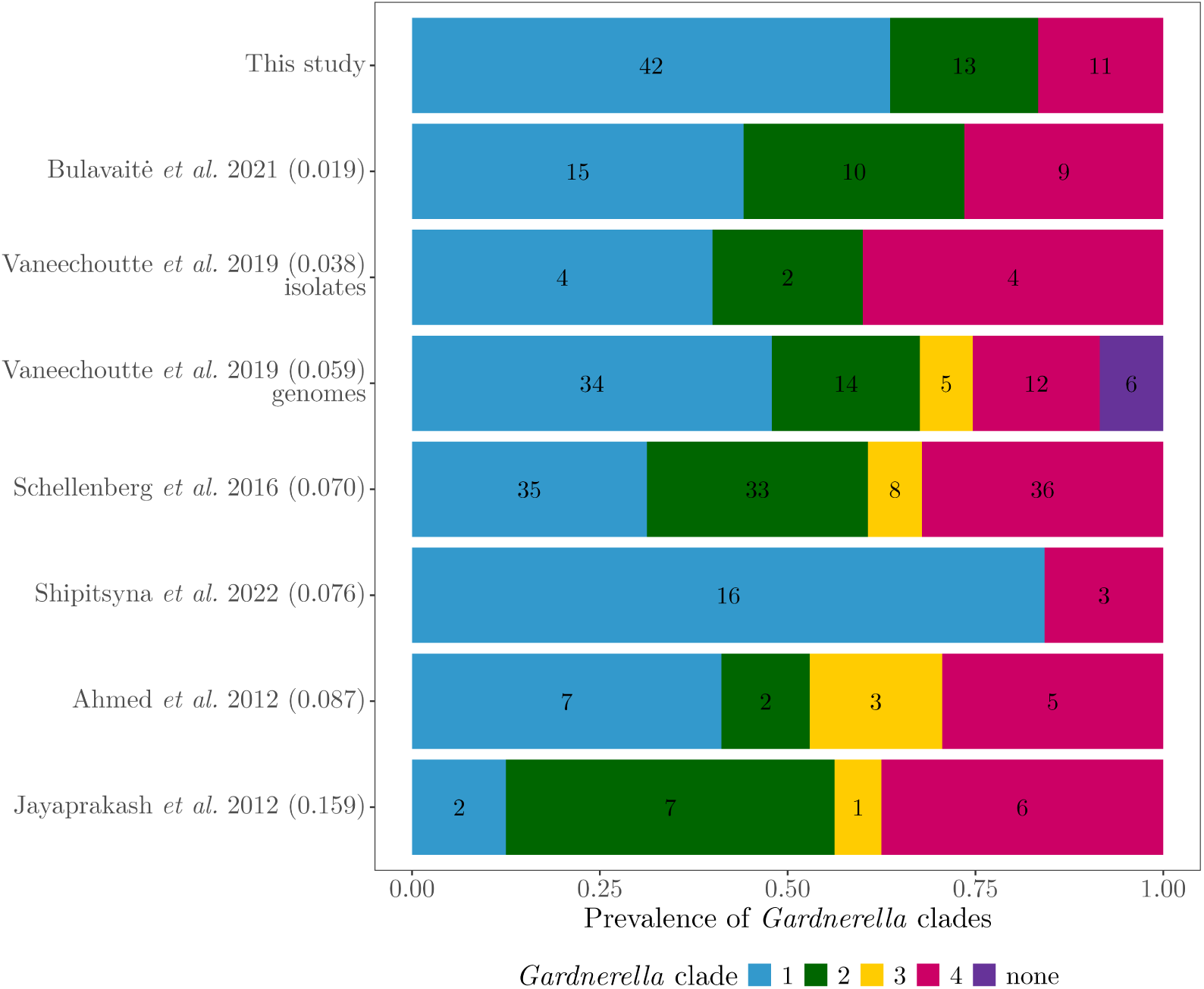
**Prevalence of *Gardnerella* clades in published datasets**. Colors show the different *Gardnerella* clades as indicated in the phylogenetic tree (Fig. 1) and numbers indicate count data for each clade in each study (shown on the y-axis). The purple clade contains isolates either not associated to the previously described four clades or associated to the fifth clade of this study. Studies are ordered from top to bottom by increasing Jensen-Shannon divergence (JSD) between our study’s clade distribution and every other study, with corresponding JSD values shown in parentheses on the y-axis. Vaneechoutte *et al.* (2019) is shown as two entries, separating study-specific clinical isolates from previously published genomes.

Our results confirm that *Gardnerella* complex can be stratified into five clades on the basis of the *cpn60* phylogenetic relationships, with clades 1, 2, and 4 being the most prevalent among our clinical isolates.

### Sialidase gene content varies across *Gardnerella* clades and diverges between *nanH* paralogs

We assessed the presence or absence of *nanH1-3* sialidase genes in clinical isolates by PCR. The most common sialidase gene was *nanH1*, present in 53/66 isolates and highly prevalent in clades 1, 2, and 3. Overall, *nanH2* was rarely detected, present in only 4/66 isolates, including *G. pickettii* and *Gardnerella* genome species 11 within clade 2. Finally, the sialidase gene *nanH3* was detected in 25/66 isolates and highly prevalent in clade 2 (present in 10/13 isolates), including *G. pickettii* and *G. piotii* (Figure 1). Classification by *cpn60* -defined clades was more consistent than expected by chance with the presence-absence patterns of the *nanH1* and *nanH3* sialidase genes, but not with *nanH2* (Fisher’s exact test values: p *<* 0.001 for *nanH1* and *nanH3* ; p = 0.28 for *nanH2* ). Although *nanH1* was detected more frequently than expected by chance in clades 1 and 2 (pairwise comparisons using Fisher’s exact test with Benjamini & Hochberg correction, all p *<* 0.001), *nanH3* was predominantly restricted to clade 2. We also observed a positive association between the presence of *nanH1* and *nanH3* (Fisher exact test: p = 0.004), whereas *nanH2* showed no association any of the other *nanH* genes (Fisher exact tests: all p = 1). Overall, taxonomic assignment within *Gardnerella* clades 1 and 2 partially predicts the presence of the *nanH1* and *nanH3* sialidase genes.

Among sialidase-related genes, *nanH1* was the most prevalent and was consistently present in clades 1, 2, and 3. We thus used *nanH1* alongside the housekeeping gene *cpn60* to investigate the evolutionary histories of our clinical isolates and the representative reference isolates described above. At the clade level, the *nanH1* and *cpn60* phylogenies were largely congruent, with clades 1, 2, and 3 forming monophyletic groups in both trees. The main discrepancy between the two phylogenies was the placement of *Gardnerella* genome species 11, which appears basal to clade 2 in the *cpn60* tree but basal to clade 3 in the *nanH1* tree (Supplementary Figure S4). The most striking difference between the two trees lies not in their overall topology, but in the distributions of branch lengths in the *nanH1* phylogeny compared to *cpn60*.

To further investigate differences in evolutionary patterns, we estimated sequence divergence within and between clades using pairwise genetic distances among isolates. Pairwise distances for *cpn60* sequences displayed a bimodal distribution (Figure 3 A, horizontal axis), with the modes largely corresponding to intraclade and the interclade distances (respectively around 0.02 and 0.12 substitutions per site). In contrast, pairwise distances in *nanH1* showed a broadly dispersed distribution, with an excess of values close to zero (Figure 3 A, vertical axis). Indeed, the distribution of pairwise distances for the two loci differed significantly (Anderson-Darling test: AD = 320.00, T.AD = 419.12, p *<* 0.01). Overall, variation in pairwise *cpn60* distances was a poor predictor of variation in pairwise *nanH1* distances (z = 5.13, p *<* 0.01, Kendall’s tau = 0.12 for intra-clade pairwise distances, Figure 3). Intraclade *nanH1* distances were relatively higher than expected given *cpn60* distances (zero-inflated gamma model and posthoc comparison between normalized intra and inter-clade *nanH1* pairwise distances, z value = -45.48, p *<* 0.01, Figure 3 B).

**Figure 3:**
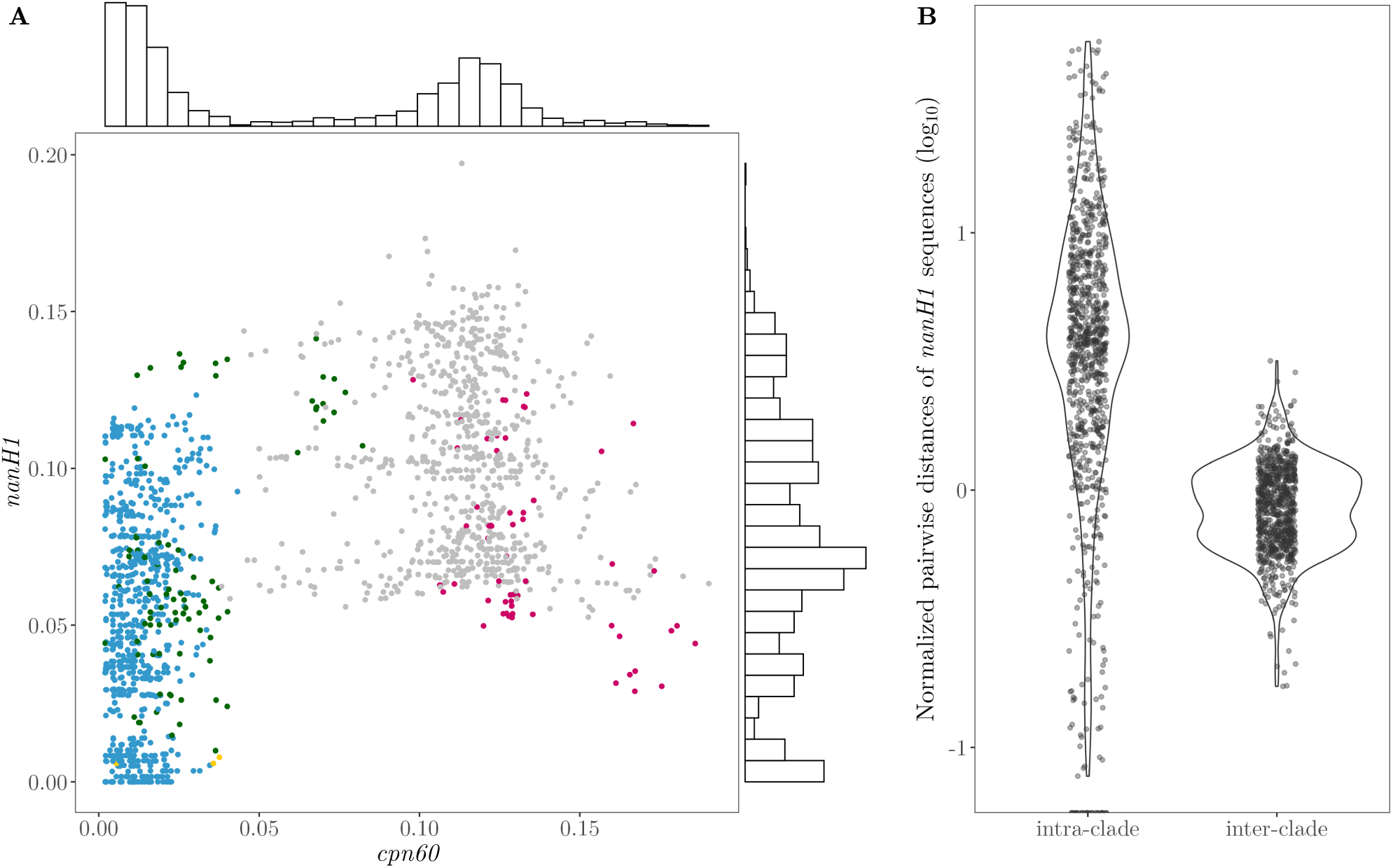
Pairwise distances in *Gardnerella* spp. between *nanH1* and *cpn60* loci. (A) Pairwise distances from DNA sequences of *cpn60* (x-axis) and *nanH1* (y-axis) from *Gardnerella* spp. clinical isolates and *Gardnerella* reference species, estimated using the Tamura and Nei (1993) model of evolution with gamma correction. Clade combinations are shown with different colors: blue, green, and yellow for pairwise distances within clades 1, 2, and 3 respectively, pink for interclade pairwise distances involving 3D6 isolate from clade 4, and grey for all other interclade distances. The distributions of pairwise distances for each gene are shown as histograms along the corresponding axis. (B) Pairwise distances from *nanH1* sequences normalized by the corresponding *cpn60* pairwise distances, within and between clades. Data are log10-transformed.

Within clade 1, *cpn60* -based pairwise distances displayed a very narrow distribution (95% CI 0.012 - 0.013), with no co-variation with the more dispersed *nanH1* -based pairwise distances (95% CI 0.05 - 0.055) (dots in blue in Figure 3 A). A similar pattern is observed within clade 2 (dots in green in Figure 3 A), with additional outliers showing larger pairwise distances corresponding to the shift in phylogenetic placement of *Gardnerella* genome species 11 between the two trees. Finally, we identified several isolates with large *cpn60* pairwise distances, corresponding to the right tail of the x-axis distribution in Figure 3 A, that nonetheless displayed closely related *nanH1* sequences (in pink). They represent between-clade pairwise comparisons involving the sole clade 4 isolate (3D6) carrying a *nanH1* gene, which notably is also the only isolate in this clade to harbour *nanH3*. Given its phylogenetic position, basal to the *nanH1* sequences of clade 2, we infer that this isolate may have acquired this sialidase gene ancestrally via horizontal transfer from an ancestor of the clade 2 lineage.

To further investigate the contrasting evolutionary patterns of *cpn60* and *nanH1*, we compared nucleotide composition at the third codon position, a sensitive indicator of codon usage bias and potential horizontal gene transfer (Figure 4). The GC-content of the two genes differed within clades 1 and 2 but not within clades 3 and 4 (interaction gene x clade Chisq(3) = 20.67, p *<* 0.001 followed Tukey-adjusted pairwise comparisons of marginal means). Overall, *cpn60* and *nanH1* GC content values were low, ranging from 0.30 for *cpn60* in clade 1 to 0.38 for *nanH1* in clade 4 (median values) Isolates in clade 1 consistently had lower GC content than those in clades 2, 3, and 4 (Chisq(3) = 316.27, p *<* 0.001 followed by Tukey-adjusted comparisons). This low overall GC content was associated an over-representation of T-ending codons in *cpn60* for all *Gardnerella* clades, whereas *nanH1* showed an over-representation of A-ending codons (interaction gene x nucleotide Chisq(3) = 7325.3, p *<* 0.001 followed Tukey-adjusted pairwise comparisons of marginal means, Figure 4 B).

**Figure 4:**
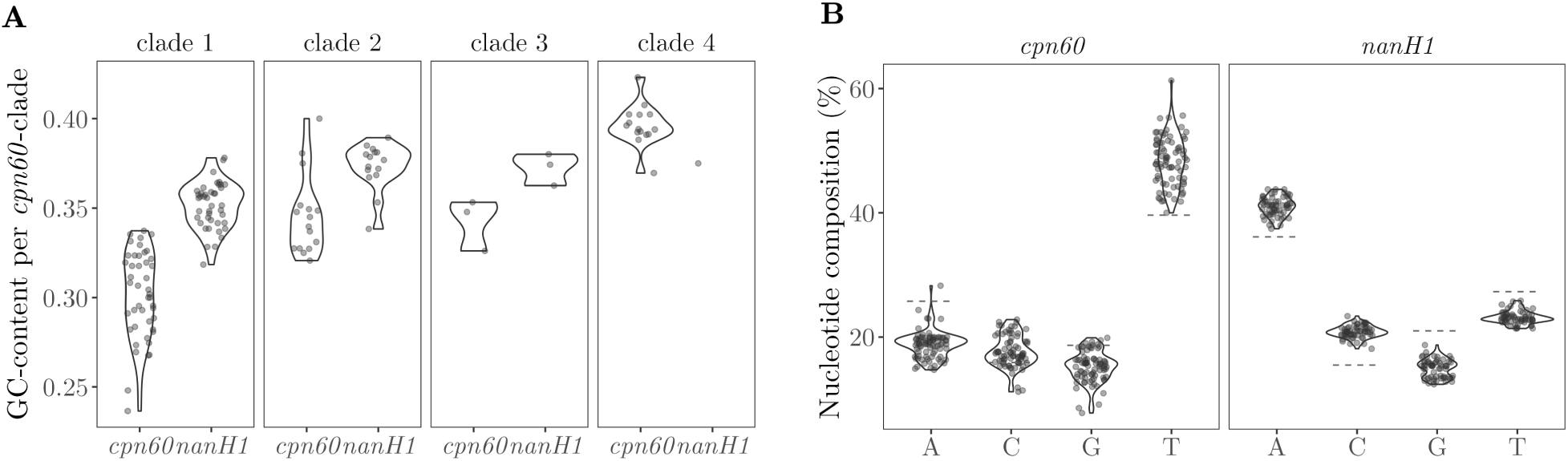
GC-content (A) and nucleotide composition (B) at third position of codon in *nanH1* and *cpn60* genes in *Gardnerella* spp. (624 and 552 bp, respectively). Values correspond to the sequences from clinical isolates communicated in this study, as well as, when available, *Gardnerella* reference genomes, obtained from the National Center for Biotechnology Information (NCBI) for *nanH1* and from the Chaperonin database (cpn60DB) for *cpn60* (Supplementary Table S6). Reference dashed lines represent median values for nucleotide frequencies at third position of codon in *Gardnerella* spp. across whole genome sequences for *cpn60*, and across sialidase genes for *nanH1*, respectively.

We conclude that the evolutionary histories of *cpn60* and of *nanH1* genes within the *Gardnerella* complex are only partially congruent. Although members of the same *Gardnerella* clade tended to display more closely related *nanH1* sequences, within clade variation of this sialidase gene was three- to five-fold greater than the corresponding variation observed for the taxonomic marker *cpn60* gene. Further, nucleotide composition of *nanH1* and *cpn60* genes differs, irrespective of taxonomy.

### High phenotypic diversity in sialidase activity within and between *Gardnerella* clades

We quantified the secreted sialidase activity for a subset of 40 *Gardnerella* clinical isolates following growth on SB solid medium. The subset was selected based on taxonomic classification, with isolates randomly sampled from each clade in proportions reflecting overall clade prevalence. We observed substantial phenotypic diversity, both qualitatively and quantitatively, encompassing sialidase-producing and non-producing isolates, as well as activity levels spanning several orders of magnitude (Figure 5, raw data in Supplementary Figure S1).

**Figure 5:**
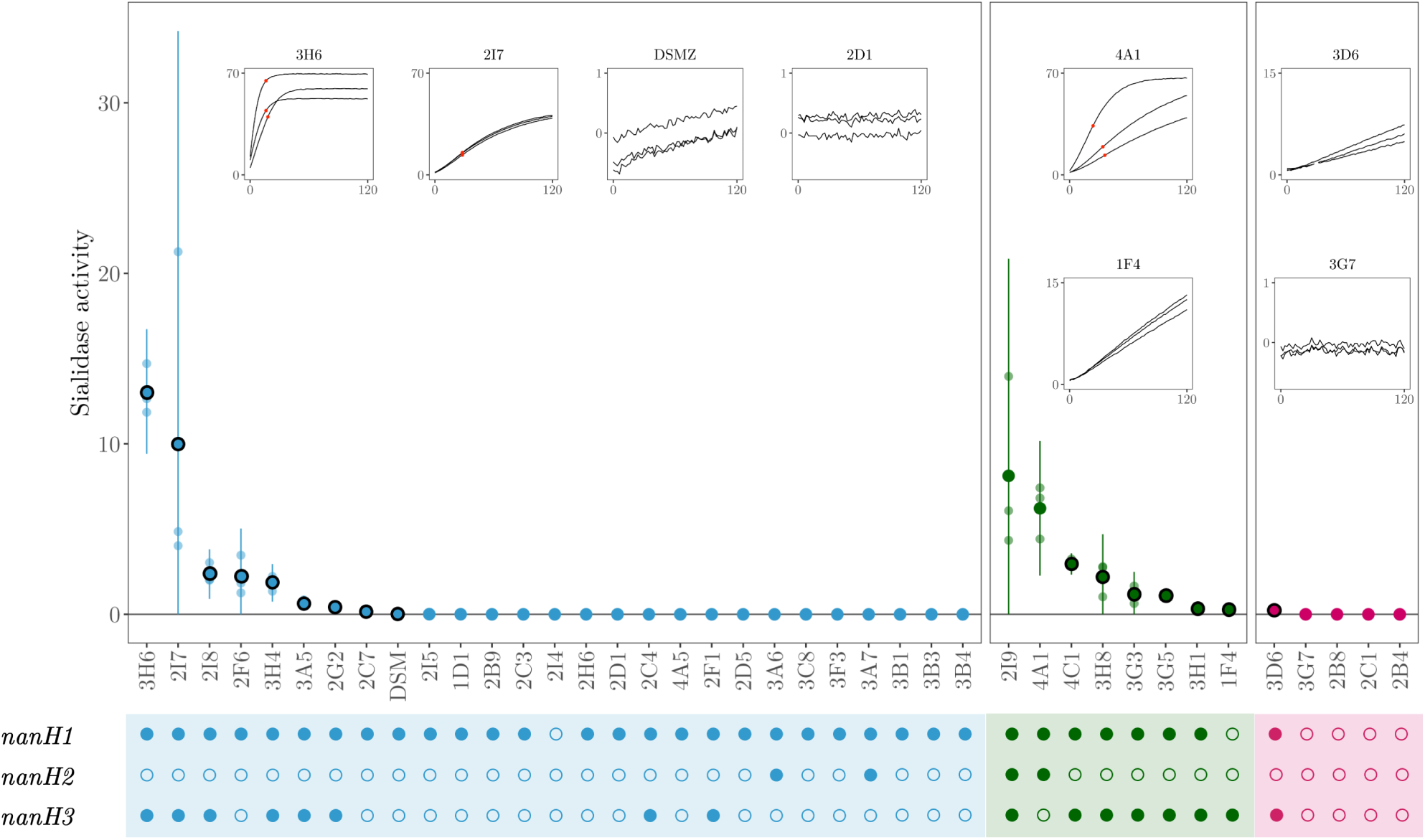
Sialidase activity of *Gardnerella* spp. clinical isolates and of the DSM *G. vaginalis* type strain. In the y-axis sialidase activity is expressed as micromols of sialic acid 4MU-sialic acid substrate hydrolysed per minute per mg/mL of secreted protein. The mean value for biological triplicates and the zero-truncated 95% confidence intervals are shown. Sialidase activities significantly greater than zero are highlighted with black circles around mean dots (linear mixed model on slope data followed by one-sample one-sided t-tests with Benjamini & Hochberg correction, p *<* 0.05). The insets show, for representative examples of assay responses, the actual raw data with the fluorescence kinetics generated by substrate hydrolysis (y-axis) over time (in minutes, x-axis). Red dots in the insets indicate the beginning of curve saturation, when applicable. The lower panel displays filled and empty circles, respectively for the presence and absence of the correspondent *nanH* sialidase gene, using the code in Figure 1.

Isolates within the same *cpn60* clade exhibited distinct sialidase activity profiles (zero-inflated gamma model, Chisq(2) = 14.56, p *<* 0.01 followed by Tukey contrasts between clades, Figure 5). All isolates from clade 2 (8/8 ) displayed detectable sialidase activity, compared with 9/27 isolates in clade 1 and 1/4 in clade 4 (clade 2 vs 4: t(114) = 3.32, p *<* 0.01; clade 1 vs 2: t(114) = 3.35, p *<* 0.01; clade 1 vs 4: t(114) = 1.06, p = 0.54). The highest sialidase producers were found within clade 1, with activity levels exceeding those of low producing isolates in the same clade by more than three orders of magnitude (a linear mixed model on slope data confirmed a strong effect of isolate identity on activity levels: Chisq(39) = 433.28, p *<* 0.001). In clade 2, six of eight isolates consistently exhibited sialidase activity, while the other two isolates were also positive but showed highly variable activity levels (isolates 2I9 and 4A1; Figure 5).

### Limited concordance between *nanH1-3* gene presence and sialidase activity in *Gardnerella*

Next, we examined the qualitative concordance between sialidase activity and presence or absence of sialidase genes among the clinical isolates. The *nanH1* gene was present in 26/27 phenotyped clinical isolates from clade 1; however, only nine of these exhibited detectable sialidase activity. After accounting for the presence of the other two sialidase genes, we found no overall association between *nanH1* presence and sialidase activity (binary logistic regression on presence / absence of sialidase activity, z value = 0.932, p = 0.35). In contrast, the presence of *nanH3* was a stronger predictor of sialidase production; it was present in 14/18 sialidase-producing isolates by in only 2/22 non-producers, corresponding to a 12-fold odds of exhibiting sialidase activity (95% CI: [4.5, 33.9], z value = 4.883, p *<* 0.001). Finally, *nanH2* was infrequently detected in our isolate collection and showed only a weak positive association with sialidase activity (95% CI: [0.9, 16.4], z value = 1.765, p = 0.078).

However, the presence or absence of the three *nanH1-3* sialidase genes, whether individually or in combination, did not fully explain the presence or absence of the sialidase-producing phenotype (Figure 5).

### Clinical BV diagnosis is not associated with *Gardnerella* taxonomy or sialidase activity

Most isolates (75%) were derived from vaginal samples obtained from women with a clinical diagnosis of BV. We aimed to test whether BV status was associated, beyond random expectations, with *Gardnerella* genotype, the isolate’s sialidase gene repertoire, and sialidase production. With respect to *Gardnerella* taxonomy, no clade was preferentially associated with clinical BV status among the 40 isolates examined (Fisher exact test for association between clade identity and BV status: p = 0.75, Figure 6). We observed a marginal association between the sialidase repertoire of the isolates and BV status (Fisher exact test: p = 0.048), likely driven by three isolates carrying only *nanH1* and *nanH2*, all of which were BV-negative (Supplementary Table S3). Finally, we found no association between clinical BV status and sialidase activity when assayed under our experimental conditions (Fisher exact test: p = 0.83).

**Figure 6:**
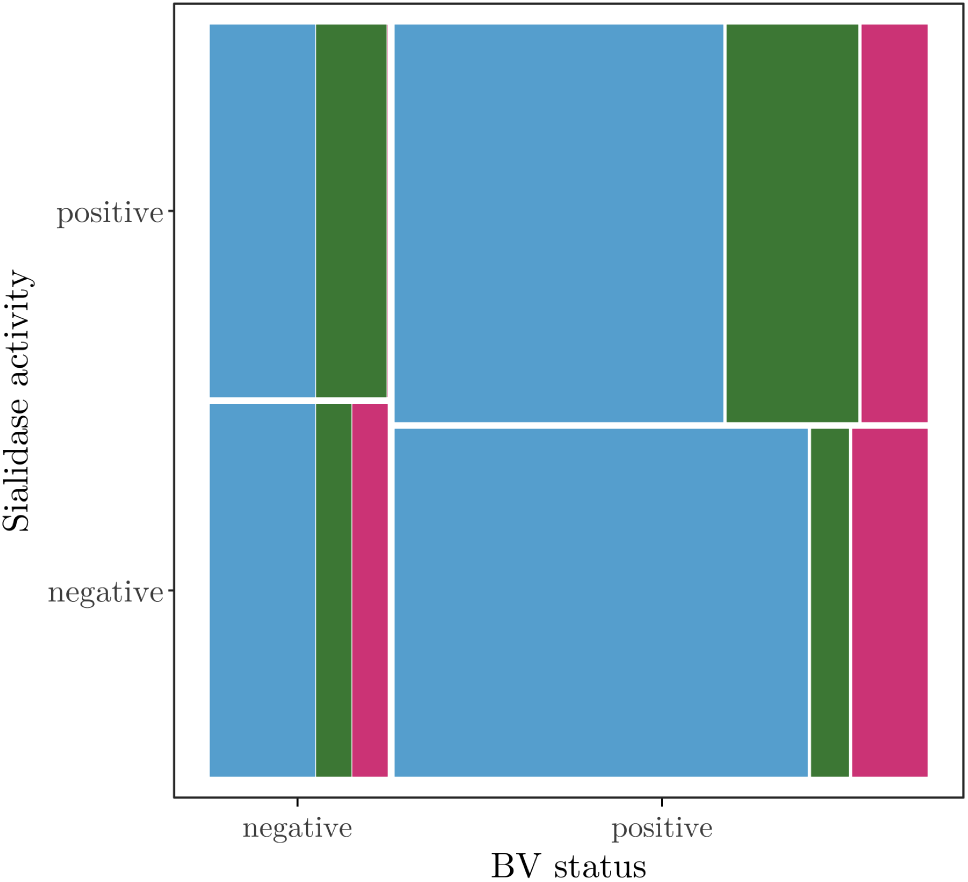
Relationship between BV diagnosis status, *Gardnerella* taxonomy, and sialidase activity. The size of each rectangle in the mosaic plot is proportional to the number of samples in the corresponding category, stratified by BV status (x-axis), detection of sialidase activity (y-axis) and *Gardnerella* taxonomy (colour code according to Figure 1, blue for clade 1, green for clade 2 and pink for clade 4).

### Substantial within- and between-clade diversity in antibiotic resistance in *Gardnerella* clinical isolates

Finally, we determined the minimum inhibitory concentration (MIC) of four antibiotics commonly used in BV treatments (metronidazole, clindamycin, moxifloxacin, and augmentin) using E-tests for 62 *Gardnerella* clinical isolates (Figure 7).

**Figure 7:**
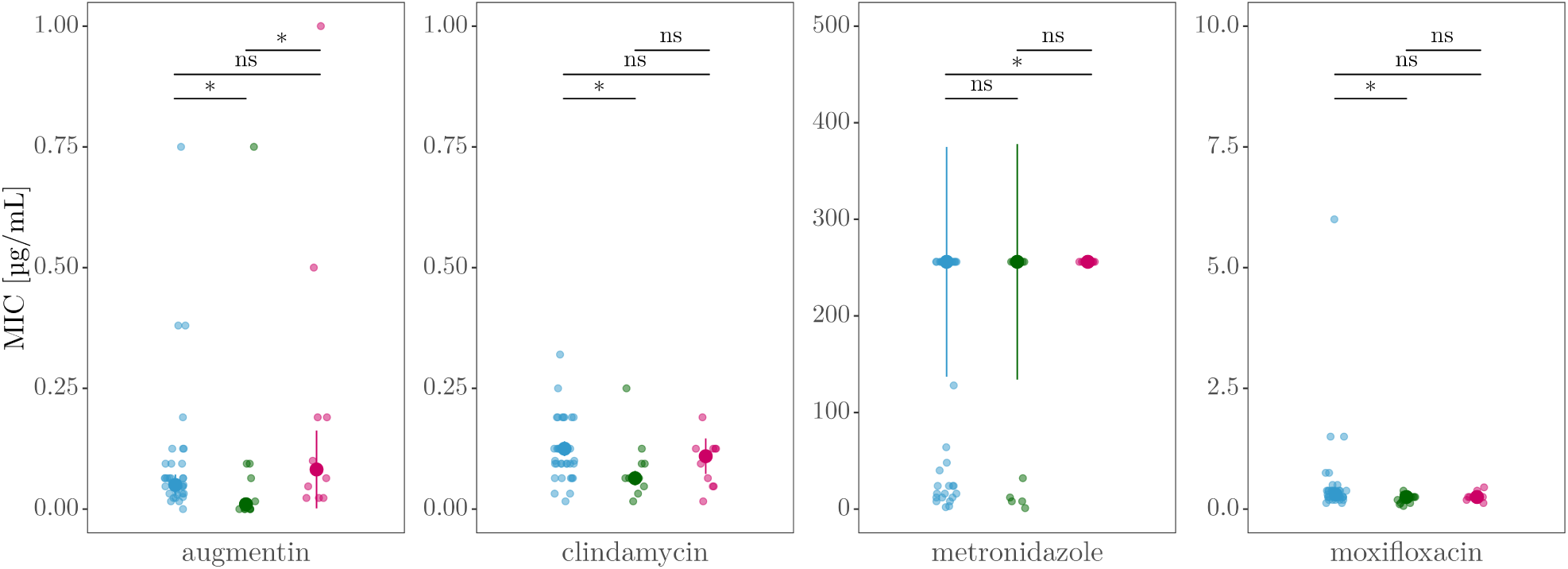
Antibiotic susceptibility of clinical *Gardnerella* isolates by taxonomic clade. Individual points represent Minimum Inhibitory Concentration (MIC) values for each isolate and antibiotic, colored by clade: blue for clade 1, green for clade 2, and pink for clade 4. Larger points indicate median MIC values, with error bars showing the interquartile range (IQR) around the median. Results of post hoc comparisons of MIC distributions between clades are indicated by asterisks for statistically significant differences (corrected *p<* 0.05) and ”ns” for non-significant results. Note the different y-axis scales across panels.

We observed clade-specific patterns of antibiotic susceptibility, with significant differences in MIC distributions among clades for all antibiotics tested (Kruskal–Wallis, corroborated by Anderson–Darling, Supplementary Table S8). Although post hoc comparisons revealed significant differences across multiple antibiotics, the most pronounced differences were observed for metronidazole. All clade 4 isolates exhibited uniformly high MICs, whereas isolates in clades 1 and 2 had bimodal susceptibility patterns (Figure 7). For augmentin, clade 2 isolates exhibited significantly lower MIC values than isolates from clades 1 and 4. Clindamycin and moxifloxacin exhibited modest but statistically significant differences in MIC values between clades 1 and 2. Finally, we did not observe clear differences in MIC distributions by BV status (Supplementary Figure S2).”

## Discussion

Despite a growing body of research, the etiological relationship between *Gardnerella* spp. and BV remains unresolved. The extensive genomic diversity within the *Gardnerella* genus likely contributes to the complexity of BV etiology. Here, we integrated taxonomic, genomic, and functional analyses to investigate the diversity in *Gardnerella* isolates derived from both BV-positive and BV-negative clinical samples. Our results reveal substantial diversity across five phylogenetic clades defined by housekeeping gene sequences, a heterogeneous *nanH* sialidase gene repertoire, and pronounced phenotypic variability in both sialidase activity and antibiotic resistance. None of these features emerged as a reliable predictor of BV clinical status.

### Most *Gardnerella* spp. isolates belong to three *cpn60* -based clades

Early efforts to stratify *Gardnerella* spp. diversity have applied a range of approaches, including neighbor-grouping analyses [41], *cpn60* -based phylogenetic subdivision [40], and genome-wide similarity metrics such as digital DNA–DNA hybridization and average nucleotide identity [43]. Despite the differences in taxonomic resolution and nomenclature, these approaches consistently recover a limited number of deeply diverged phylogenetic lineages. Consistent with this convergence, we found that the 13 *Gardnerella* species delineated by Vaneechoutte and colleagues [43] collapse into five robust phylogenetic groups based on *cpn60* sequence variation, four of which correspond to previously described clades (Table 1). The congruence between these classifications supports the robustness of the underlying phylogenetic structure and confirms that *cpn60* gene sequences are excellent predictors of whole-genome relationships in *Gardnerella* spp. [40, 51].

Most of our clinical isolates belong to clades 1, 2, and 4, with clade 1 being predominant and the majority of isolates corresponding to *G. vaginalis*–like lineages. This pattern is consistent with other reports identifying these clades as dominant among vaginal isolates from women of reproductive age [29, 43], although some studies have reported more balanced [25] or divergent distributions [40, 41]. Such variation among studies may reflect ecological and geographic differences, as well as methodological differences in isolate sampling and taxonomic assignment. More recently, several studies have proposed revised phylogenetic frameworks for the *Gardnerella* complex [52–54], with one preprint proposing an alternative nomenclature for the genus [55].

In our study, each taxonomic clade is characterized by low genomic diversity, reflected in the relatively low intra-clade *cpn60* pairwise distances among our isolates. While this might suggest that most isolates fall into one of three broad categories (clades 1,2 and 4), our sialidase gene–related data reveal a more complex picture.

### Sialidase *nanH* genes repertoire in clinical isolates does not match *Gardnerella* taxonomy

Taxonomic identification alone does not accurately predict the precise combination of *nanH* genes present in a *Gardnerella* isolate. In our collection, *nanH1* was present in most genomes from clades 1, 2, and 3, while *nanH2* is only rarely detected across any clade, consistent with previous studies (Supplementary Table **??**). In contrast, *nanH3* is primarily associated with clade 2 and a subset of clade 1 isolates. As a result of these distributions, isolates within the most prevalent clades (1, 2, and 4) display a highly diverse *nanH* gene repertoire. In clade 1 and 2, some isolates possess only *nanH1*, others carry *nanH1* and *nanH3*, a few harbor *nanH1* and *nanH2*, and only two isolates from clade 2 contain all three *nanH* genes. In clade 4, most isolates lack *nanH* genes altogether, with the single exception of one isolate carrying both *nanH1* and *nanH3*.

The restricted distribution of *nanH3* raises the possibility that the gene has either been lost over evolutionary time or acquired through horizontal gene transfer as has already been suggested previously [30].

The high prevalence of *nanH1* in our collection allows comparison of its phylogenetic relationships with those inferred for the conserved *cpn60* gene. While phylogetic inferences are broadly congruent with one another (all three clades remain monophyletic), two lines of evidence suggest that the evolutionary trajectories of the two loci are only partly linked. Indeed, branch length patterns differ: *cpn60* pairwise distances are bimodal and reflect phylogeny, while *nanH1* pairwise distances are overdispersed and not informed by taxonomy (Fig. 4A). Accordingly, intraclade pairwise distances at the *nanH1* locus are a median 3.8-fold higher than those observed for *cpn60*, with substantial variation among isolate pairs. Further, clade 2 is well defined and supported in the *cpn60* phylogeny, with *Gardnerella* genome species 11 positioned basally within the clade. In contrast, clade 2 is less clearly defined in the *nanH1* phylogeny, where *Gardnerella* genome species 11 appears as basal to clade 3. Finally, isolate 3D6 is exceptional among clade 4 members, not only because it carries both *nanH1* and *nanH3*, but also because it appears basal to clade 2 in the *nanH1* phylogeny. The characteristics of this isolate are consistent with horizontal transfer events involving both *nanH* genes. Molecular compositional signatures further support this hypothesis: GC content and third-codon-position nucleotide composition differ between *nanH1* and *cpn60*, consistent with acquisition from donors with distinct mutational biases and nucleotide composition.

### From genotype to phenotype: *nanH* genes repertoire and sialidase activity

The presence of a gene reflects potential function only; its expression depends on regulatory and ecological context. Historically, sialidase production in *Gardnerella* spp. was attributed primarily to *nanH1* [23, 26, 27].

However, this monogenic view has been challenged by the identification of *nanH2* and *nanH3* as additional contributors to sialidase-producing phenotypes [28]. The decoupling we observed between *nanH1* presence and secreted sialidase activity is consistent with these recent findings, with *nanH3* emerging as the strongest predictor of sialidase activity in our isolates.

The biochemical conditions of our enzyme assay may have contributed to the observed mismatch between gene presence and enzyme activity. We have quantified secreted sialidase activity by resuspending bacterial colonies grown on solid medium, which primarily captures secreted enzyme activity and may not detect cell-associated sialidase. Proposed differences in enzyme localization (*nanH2 and nanH3* are thought to be largely extracellular [28], whereas *nanH1* has been suggested to be intracellular [30, 39]) may therefore contribute to the stronger association between sialidase activity and *nanH3* in our supernatant-based assay. However, previous studies have shown that isolates negative in supernatant-based fluorometric assays were also negative in filter spot assays performed with whole cultures [29] and that a significant fraction of *Gardnerella* sialidase activity can be found in culture supernatants [23, 28]. Furthermore, enzyme activity measured *in vitro* using artificial substrates may not accurately reflect *in vivo* activity on natural substrates or within the complex biochemical environment of the host. In our case, we used whole culture supernatants without protein purification. It is also possible that bacterial sialidase inhibitors present in the supernatant suppressed detectable activity, even when the enzyme was present. Finally, the assay substrate consists of a sialic acid moiety linked to a fluorogenic 4-methylumbelliferone group, which does not chemically resemble the sialylated glycans found in the vaginal epithelium. Although substrate cleavage requires sialidase activity, limiting false positives, the biochemical conditions in the supernatant may allow enzymes that efficiently cleave natural substrates to show little or no activity against the 4-methylumbelliferone–sialic acid substrate, potentially resulting in false negatives.

Other studies measuring sialidase activity directly from bacterial cultures have similarly reported that *nanH1* is not predictive of activity [25, 27–29, 31] and presented stronger evidence for *nanH3* [28]. Preliminary reports suggest that *nanH3* is directly linked to sialidase production, as gene ablation results in loss of secreted sialidase activity and mucus degradation in *G. pickettii* [56]. Furthermore, high cervicovaginal *nanH3* gene load has been associated with BV and increased persistence of high-risk HPV16 infection [57]. In our study, the presence of *nanH3* was associated with sialidase activity, but it did not fully account for the phenotypic diversity observed. Similar reports of discordance between gene content and sialidase expression in *Gardnerella* spp. highlight the likely role of regulatory and environmental factors in shaping phenotypic diversity [29–31].

It remains possible that the different *nanH* genes are not functionally redundant. They may differ in substrate specificity, regulation, timing of expression, or ecological function, potentially allowing specific strains to adapt to different host conditions. For instance, some sialidases may be constitutively expressed, while others are inducible under host- or microbiota-derived cues, or they may differ in substrate specificity. Elucidating such functional differentiation, for example through heterologous expression, targeted mutagenesis, and biochemical characterization of the corresponding enzymes, will help to understand how sialidase-mediated interactions contribute to microbial ecology and BV pathology. Proteomic analyses of culture supernatants and vaginal secretions could identify which sialidases are secreted under physiologically relevant conditions, and the cues that regulate their expression. These data would inform the development of targeted therapeutic strategies aimed at selectively inhibiting specific sialidase variants.

Our results emphasize the limitations of inferring microbiota phenotype and clinical relevance solely from taxonomic or genomic information. Functional assays remain essential and must be interpreted in the light of environmental conditions, assay design, and regulatory complexity.

### Taxonomy and antibiotic resistance profile

The complex genotype-phenotype relationship observed for sialidase activity extended to bacterial taxonomy and antibiotic resistance. We detected differences in minimum inhibitory concentrations between clades for all four antibiotics tested, although no obvious clade-specific resistance profile could be defined. Metronidazole was distinctive: all clade 4 isolates were highly resistant whereas clade 1 and 2 isolates displayed bimodal distributions, comprising both resistant and sensitive isolates. This pattern is consistent with findings from at least two other studies, which reported bimodal metronidazole resistance phenotypes in clade 1 and 2, but uniformly resistant isolates in clade 3 and 4 (MIC ≥ 32 µg/mL) [37, 54]. For other antibiotics, resistance patterns were less clearly tied to taxonomy. Clade 2 isolates tended to be more susceptible to augmentin than clades 1 and 4, with modest but significant clade-associated differences also detected for clindamycin and moxifloxacin. Overall, clade identity may reflect broad trends in antibiotic susceptibility, but it cannot serve as a reliable predictor of resistance at the individual isolate level.

### Quantitative sialidase activity reveals extensive functional heterogeneity across the *Gardnerella* complex

We observed a marked disconnect between the phylogenetic structure of *Gardnerella* spp. and the distribution of key phenotypic traits, with sialidase activity, antibiotic resistance, and clinical associations with BV largely unconstrained by clade boundaries. Sialidase activity, in particular, exhibited substantial quantitative variation both between and within clades.

Although all clade 2 isolates produced detectable sialidase activity, the strongest producers in our dataset originated from clade 1. Clade 1 isolates displayed pronounced phenotypic heterogeneity, ranging from high producers to complete non-producers despite widespread carriage of *nanH1*. These patterns are consistent with previous reports describing uniformly sialidase-positive clade 2 isolates and variable activity within clade 1 [30, 31, 39, 58] (see Supplementary Table S9). However, our taxonomic resolution does not allow species-level assignment of all isolates, limiting our ability to determine whether high-producing clade 1 isolates predominantly correspond to G. vaginalis, which has been reported to include sialidase-producing strains [39], while non-producing clade 1 isolates correspond mainly to genome species 2, currently characterized as non-producing [39]. Similarly, the species affiliation of isolate 3D6 remains uncertain, which, to our knowledge, represents the first clade 4 isolate reported to exhibit sialidase activity.

Finally, we detected low but reproducible sialidase activity in the *G. vaginalis* type strain DSM 4944 (=ATCC 14018), which has previously been reported as a non-producer [43]. Together, these observations highlight the limitations of categorical classifications of sialidase activity and underscore the value of quantitative measurements for capturing functional diversity across the Gardnerella complex.

### Ecological and clinical relevance

The absence of strong associations between *Gardnerella* genotypes or phenotypes and clinical BV highlights the need for a systems-level understanding of bacterial communities and disease risk. It suggests that the contribution of *Gardnerella* lineages to BV may not reside in taxonomic identity or individual virulence traits alone, but rather in the ecological context of the vaginal microbiota, including interactions with other microbes [59] and host factors [60, 61]. Indeed, a recent study has shown that no *Gardnerella* species was a specific marker for BV and hypothesized that BV reflects the presence of high *Gardnerella* species diversity [62]. Furthermore, other BV-associated bacteria, such as *P. bivia* or *Fannyhessea vaginae* (formerly known as *Atopobium vaginae*, have long been identified as key players in the induction of BV [5], highlighting the need for more complex community assays.

Although our study includes a relatively large number of cultured *Gardnerella* clinical isolates, it is enriched for BV-positive samples, which may limit power to detect associations with BV status. Moreover, analysis of a single isolate per woman may underestimate within-host diversity. Nevertheless, prior work indicates that vaginal microbiota are typically dominated by a single *Gardnerella* lineage despite frequent co-occurrence of multiple species [51], suggesting that our isolates capture the dominant population in most samples.

Our assay measured sialidase activity from colonies grown in isolation on rich solid medium, conditions that may not recapitulate *in vivo* expression patterns. *Gardnerella* spp. may alter their secretion repertoire when growing within a multispecies community, under lower nutrient conditions, or in response to host signals such as pH, mucus components, or immune factors. As a result, strains that appear inactive under laboratory conditions may still produce sialidase in the vaginal environment, while high *in vitro* activity may not translate directly to *in vivo* function. From a clinical perspective, this reinforces the need for integrative diagnostics that go beyond single-marker tests and consider ecological and functional dimensions of microbial communities.

Collectively, these findings call into question the clinical interpretability of sialidase-based diagnostics, such as the chromogenic “blue test” used in some over-the-counter BV kits. Our data do not support a direct link between sialidase activity and BV status, suggesting that sialidase detection alone may be an unreliable surrogate for BV. Our results also do not support earlier suggestions of the potential use of *nanH2* or *nanH3* as molecular diagnostic markers of BV [28, 57]. More informative sampling would combine microbiological characterization, clinical diagnosis, and quantitative sialidase testing (eg. luciferase-based methods) within the same specimens, helping to distinguish when sialidase activity reflects physiologically relevant processes and when it does not.

## Conclusion

We present a paradigmatic example of discordance between genetic content, taxonomic stratification, biochemical function, and clinical phenotype in *Gardnerella* species. Our findings emphasize how limited our current ability remains to infer microbial community contributions from the genomic data of individual members alone. As our understanding about bacterial genomics grows, integrating functional data is essential to avoid shortcuts in interpreting biochemical activity or clinical relevance. Our results for *Gardnerella* and BV showcase how historical contingency, horizontal gene transfer, and strain- and context-specific expression patterns shape phenotypic diversity, beyond taxonomy and gene annotation. Our results highlight the sustained need for integrating physiology, plasticity, and ecology to understand the emergence and significance of microbial traits in the real world.

## Methods

### Bacterial isolates and culture conditions

Clinical isolates of *Gardnerella* spp. were obtained from vaginal samples at the Arnaud de Villeneuve Hospital in Montpellier, France (approval number from the Institutional Review Board of Montpellier University Hospital: IRB-MTP 2021 10 202100901). Vaginal swabs were streaked onto Columbia Naladixic Acid agar supplemented with 5% sheep blood (CNA, BioMérieux, Marcy-l’Étoile, France) and grown at 35°C in air-tight containers with anaerobic atmosphere generation bags (#68061; Sigma-Aldrich, St. Louis, USA). The identity of isolated colonies was verified with Matrix Assisted Laser Desorption Ionization - Time of Flight (MALDI-TOF) mass spectrometry (Bruker Daltonics, Billerica, USA). Colonies of *Gardnerella* spp. (one colony per vaginal sample) were then re-streaked onto Columbia agar (BioMérieux) at least three times and their identity confirmed each time with MALDI-TOF mass spectrometry. Freezer stocks of *Gardnerella* spp. were made by harvesting all bacterial material from Columbia agar plates and resuspending it in phosphate-buffered saline (PBS: 0.137 M sodium chloride NaCl, 2.7 mM potassium chloride KCl, 10 mM sodium phosphate dibasic Na2HPO4, 1.8 mM potassium phosphate monobasic KH2PO4; pH = 7.4) with 15% glycerol. All samples and back up aliquots were stored at -80 °C. Each vaginal sample was scored based on Spiegel’s criteria [63] for BV: ”1” indicating predominance of *Lactobacilli*, ”2” indicating the presence of *Gardnerella* spp. morphotypes together with *Lactobacilli*, and ”3” indicating rare *Lactobacilli* and predominance of other morphotypes. Samples with score of 3 are considered indicative of BV. The type strain of *G. vaginalis* (DSM 4944) was obtained from the Deutsche Sammlung von Mikroorganismen und Zellkulturen (DSMZ) and -80 °C freezer stocks were prepared as described above. A total of 66 clinical isolates were obtained between 2015 and 2016, including 48 from BV-positive samples, 17 from BV-negative samples and 1 from a sample of unknown BV status.

All isolates were recovered from freezer stocks for experimental procedures on Schaedler medium supplemented with 5% (v/v) defibrinated sheep blood (SB; SR0051E from Thermo Fisher Scientific, Waltham, USA) and 1.2% agar (SB agar). Plates were incubated in air-tight container with an anaerobic atmosphere generation bag (Sigma Aldrich, St. Louis, USA) and incubated at 37 °C for three days.

### DNA extraction and PCR amplifications

We targeted three sialidase-related genes: *nanH1*, *nanH2*, and *nanH3* [28], as well as *cpn60* gene for species identification, as *cpn60* sequences were shown to be “excellent predictors of whole genome sequence relationships” [40] and can resolve the newly described *Gardnerella* species [51].

DNA was extracted from bacterial colonies using InstaGene matrix (Bio-Rad, Hercules, USA), according to the manufacturer’s instructions. PCR conditions and mixes are described in Supplementary Tables S4 and S5. Successful amplifications of *cpn60* (552 bp) and sialidase (*nanH1* 682 bp, *nanH2* 348 bp, *nanH3* 322 bp) genes were sequenced using Sanger protocol (Genoscreen, Lille, France or Eurofins Genomics, Ebersberg, Germany).

### Sequence analysis and phylogeny inference

The quality of the *cpn60* gene sequence was evaluated using Geneious version 10.2.3, (Biomatters, https://www.geneious.com) and sialidase gene sequences were evaluated using CodonCode Aligner version 10.0.1 (CodonCode Corporation, https://www.codoncode.com). Consensus sequences from forward and reverse reads of Sanger sequencing of each strain were assembled, and mismatches or unassigned nucleotides were corrected, where appropriate, based on the electropherograms of both sequences. *cnp60* and sialidase gene sequences from reference strains of *Gardnerella* spp. were retrieved from the Chaperonin database (cpn60DB [64]) and from the National Center for Biotechnology Information (NCBI [65]), respectively, either as directly annotated and available sequences or after a blast search on all available contigs (Supplementary Table S6).

Both *cpn60* and *nanH1* sequences were aligned with MUSCLE v5 [66] at nucleotide-codon level. The *cpn60* alignment contained 554 sites, 366 patterns, 14. 3% gaps, and 72. 0% invariant sites. The *nanH1* alignment contained 625 sites, 230 patterns, 4.57% gaps, and 70.40% invariant sites. Nucleotide composition at the third position of codon was calculated in MEGA 12 software [67]. Phylogenetic inference was performed at the nucleotide level with RAxML-NG v.1.1.0[68] , using the GTR+F+gamma evolutionary model, using 10 random plus 10 parsimony start trees and performing 1,000 non-parametric sequence bootstraps.

### Measurement of sialidase activity

We modified existing sialidase activity protocols [23, 25, 28] in two main ways: (i) we measured the activity from bacterial cultures on spatially structured medium, and (ii) we standardized our measurements by total protein concentration. Spatially structured media provide a more realistic model of the type of environmental structure in which sialidase production occurs in the vaginal environment. Normalizing to total protein production ensured that negative results do not reflect inefficient protein extraction, and allowed comparison of each strain’s contribution to sialidase activity.

We measured sialidase activity, as illustrated in Figure 8, of 40 taxonomically guided *Gardnerella* spp. isolates (27 in clade 1, 8 in clade 2, and 5 in clade 4 out of 40 isolates total). All bacterial isolates were grown from freezer stocks as described above. After three days of anaerobic growth, a loopful of each strain was resuspended in 500 µL acetate buffer (0.15 M sodium chloride NaCl, 0.004 M calcium chloride CaCl2, 0.1 M sodium acetate C2H3NaO2; pH = 5.5) and adjusted to an optical density of 1 (OD_600_) using the FLUOstar Omega spectrometer (BMG LabTech, Ortenberg, Germany). 20 µL of each adjusted culture were inoculated onto sterile 2.25 cm^2^ paper squares (separation paper from MF-Millipore MCE membrane filters, Sigma-Aldrich), previously soaked in SB liquid medium and placed onto 5 mL of SB agar in 6-well tissue plates (Sigma-Aldrich), in triplicate. The experiment was carried out in blocks, each of which included two blank samples of 20 µL acetate buffer. The inoculated plates were incubated right-side up at 37 °C under anaerobic conditions for three days, after which we recovered the bacterial cultures by transferring the paper squares into 500 µL acetate buffer in 1.5 mL conical microcentrifuge tubes (Starlab). The tubes were vortexed and sonicated with a Bioruptor® Pico sonication device (Diagenode SA, Seraing Ougrée, Belgium) for two cycles of 30 seconds to remove bacteria from paper squares. After sonication, bacterial resuspensions were adjusted to an OD_600_ of 0.8 with acetate buffer, whenever possible (see Supplementary Table S7 for details of replicates with lower OD_600_), and culture supernatants were used for sialidase activity quantification.

**Figure 8:**
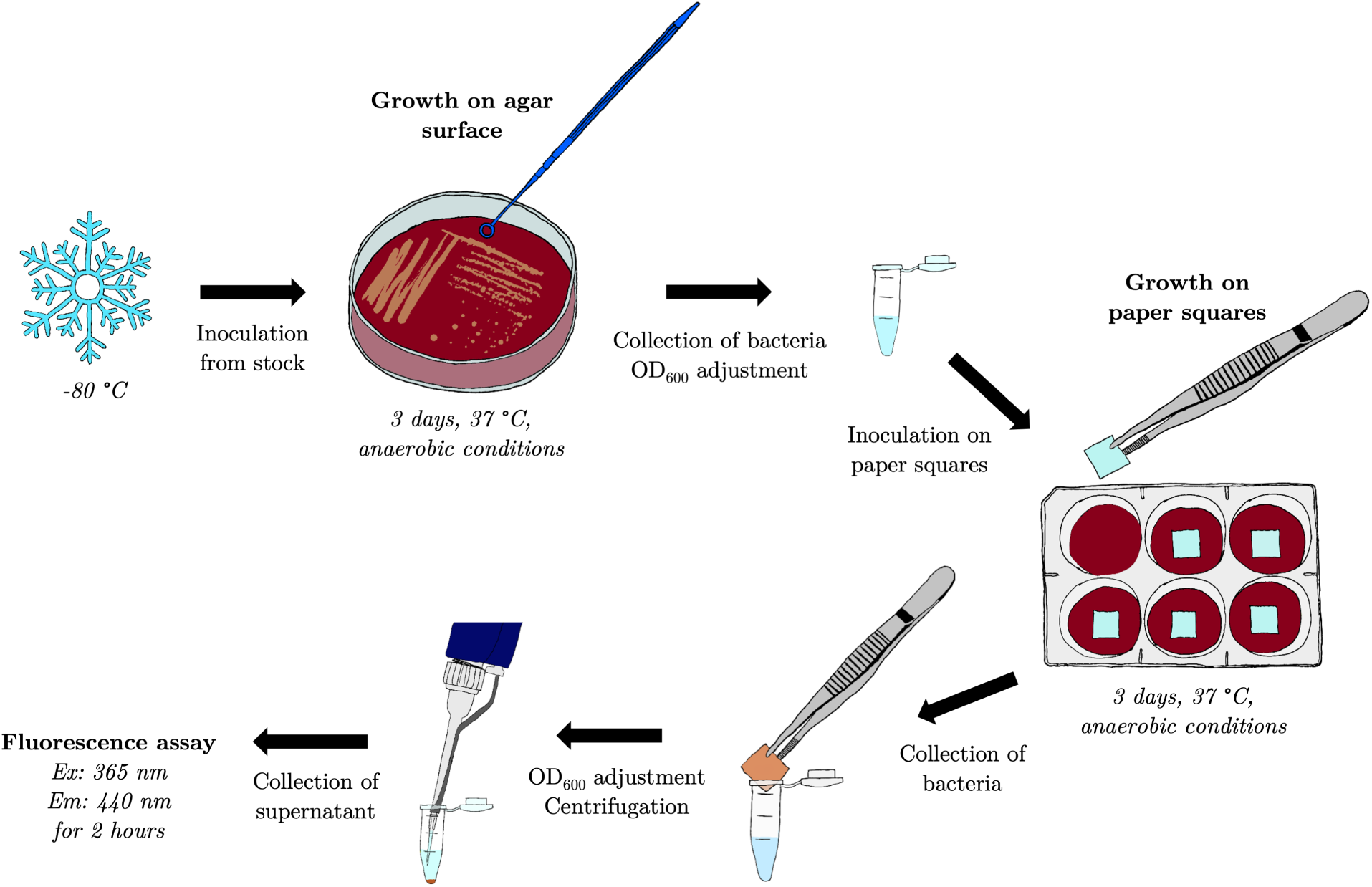
Experimental procedures for sialidase activity quantification. Bacterial isolates were inoculated from freezer stock on SB agar and incubated at 37 °C under anaerobic conditions. After three days, the cultures were recovered from agar plates and we adjusted their densities to an OD_600_ of 1. Adjusted cultures were then inoculated onto paper squares placed on SB agar in 6-well plates and incubated at 37 °C under anaerobic conditions. Three days later, cultures were collected from paper squares in acetate buffer by vortexing and sonicating. We adjusted the resuspended cultures to an OD_600_ of 0.8 and centrifuged them to isolate supernatants, which were subsequently used in sialidase activity fluorimetric assays.

Sialidase activity in culture supernatants was quantified by monitoring hydrolysis of 2^′^-(4-methylumbelliferyl) -*α*-D-N-acetylneuraminic acid sodium salt hydrate (4MU–sialic acid; Sigma-Aldrich, M8639), measured as accumulation of the fluorescent product 4-methylumbelliferone (4MU). All sialidase activity measurements were performed in Greiner CELLSTAR® 96-well flat bottom black polystyrene plates with micro-clear bottom (Sigma-Aldrich). For each sample, 50 µL of adjusted supernatant was incubated for two hours at 37 °C with 100 µL acetate buffer containing 100 µM 4MU-sialic acid. Fluorescence (excitation 360 nm, emission 430 nm) was measured with the FLUOstar Omega fluorometer every two minutes after shaking for 5 seconds at 300 rpm in a double orbital movement, and measurements were carried out with bottom optic and spiral averaging of 20 flashes per well. In every plate, we included one positive control and two negative controls (each in triplicate). The positive control contained 0.125 mU of commercial sialidase from *Clostridium perfringens* (*C. welchii* ) (N2876 from Sigma-Aldrich) in 150 µL acetate buffer with 100 µM 4MU-sialic acid. The first negative control corresponds to our blank samples and consisted of 50 µL of supernatant from one of the blank samples grown on paper squares. The second negative control was a 50 µL mixture of supernatants from three bacterial samples that we boiled at 100 °C for 10 minutes to denature sialidase enzymes. We added 100 µL acetate buffer containing 100 µM 4MU-sialic acid to both negative controls. A standard curve of 4MU (Sigma-Aldrich, M1381) at 0, 2, 5, 10, 20, and 50 µM was run in parallel on each plate to quantify the rate of 4MU–sialic acid hydrolysis. We ran duplicate sialidase activity measurements for all samples, including standards and controls.

### Total protein concentration quantification

To estimate total protein concentration in culture supernatants, we ran Bradford assays for all samples and negative controls with the same experimental block design as in the sialidase activity assays. All samples were run in 96-well plates together with a standard curve of bovine serum albumin (BSA) with concentrations of 0, 0.01, 0.025, 0.05, 0.1, 0.25 and 0.5 mg/mL. We mixed 10 µL of supernatant or standard with 200 µL ambient temperature Bradford reagent (B6916 from Sigma-Aldrich) and incubated the plates at room temperature for 30 min, before measuring absorbance (OD_600_) in the spectrometer. All samples, standards, and controls were run in duplicates.

### Estimation of sialidase activity as normalized 4MU-sialic acid hydrolysation

Sialidase activity was estimated from the rate of 4MU–sialic acid hydrolysis, calculated as the slope of the fluorescence curve during the assay. Kinetic data were smoothed using a triangular moving average with a five–data point window, after which rolling slopes were calculated using the same window size. For high sialidase–producing samples, this approach likely underestimates peak activity because the rolling window excludes the earliest, steepest increases; however, the same slope calculation was applied uniformly across all samples for consistency. To account for highly different kinetic profiles across samples, we calculated the mean of the rolling slopes across all time points before saturation. The beginning of the saturation is determined as first time point for which the slope is decreasing for 15 consecutive time points (shown as red dots in Supplementary Figure S1). This approach reduces bias from signal saturation at later time points, when the 4MU–sialic acid substrate becomes depleted. For isolates with low sialidase activity that did not reach saturation, slopes were averaged across the full time course to reduce temporal variability. Slopes were then normalized to total protein concentration, and sialidase activity was reported as µM 4MU–sialic acid hydrolyzed per minute per mg/mL of total protein.

### Antibiotic sensitivity quantification

We quantified the antibiotic sensitivity of each bacterial isolate using E-tests (Liofilchem MIC test strip, Italy), under anaerobic environments. We first streaked each isolate onto Columbia blood agar plates and incubated them anaerobically for 48 hours. We then resuspended a single colony in sterile water to reach a turbidity of 1.0 McFarland, measured with a spectrophotometer at 625 nm. We evenly spread the bacterial suspension in three directions on fresh Columbia blood agar plates before we applied E-test strips containing either Amoxicillin+Clavulanic acid (Augmentin, AUG, #92024), Clindamycin (CD, #92072), Metronidazole (MTZ, #92087), or Moxifloxacin (MXF, #92090) in the middle of the agar surface. After 48 hours of incubation under anaerobic conditions, we recorded the MIC based on the zones of growth inhibition.

A complete overview of isolate-level characteristics, including BV status, taxonomy, *nanH* gene content, sialidase activity, and antibiotic susceptibility, is provided in Supplementary Table S1.

### Statistical analysis

All statistical analyses were performed using R version 4.1.0 [69] and Rstudio version 2022.12.0+353 [70].

To analyse the distribution of isolates and *nanH* genes in *cpn60* clades, we used Fisher’s Exact Tests, with multiple comparison adjustment using Benjamini-Hochberg (BH) correction, in the RVAideMemoire package [71] . To compare the prevalence of *Gardnerella* clades among our clinical isolates with those of other studies, we searched the literature for publications that include a classification of *Gardnerella* clinical isolates similar to that of Jayaprakash *et al.* (subgroups)[40], Ahmed *et al.* (clades)[41], and Vaneechoutte *et al.* (species) [43]. We found six studies, published between 2012 and 2022, that included a *Gardnerella* spp. classification with sample sizes ranging from 10 to 112 isolates. We only considered data of clinical isolates from each study and excluded data retrieved from published genomes to avoid multiple appearances of the same sequences in our dataset. As a seventh study, we added the most recent study with a comprehensive analysis of published *Gardnerella* genome sequences (71 genome sequences in Vaneechoutte *et al.* (2019)[43]). We tested for statistical difference between the clade distribution of our study and those of each of the seven studies found in the literature by conducting multinomial tests (EMT package [72]) for all six comparisons, with Benjamini-Hochberg (BH) multiple comparison correction.

We used a zero-inflated gamma model, as implemented in the glmmTMB package [73, 74], to analyse the link between sialidase activity (as continuous quantitative variable) and *cpn60* clades, using the identity of the isolates nested into the experimental plate identity as random factor. Downstream analyses were performed using DHARMa [75] package for model evaluation and validation simulations, and car [76] and emmeans [77] packages for analysis of variance and contrasts on estimated marginal means. To identify which isolates display positive sialidase activity, we first ran a linear mixed model on slope data using the identity of the isolates as fixed effect and the experimental plate identity as random factor; and then compared each isolate slope measurements to zero with one-sample one-sided t-tests, with Benjamini-Hochberg (BH) correction for multiple testing. We used the output of these tests to create a binary variable (positive or negative) for sialidase activity. Due to between-replicate variability, slope estimates for isolates 2I9 and 4A1 were not significantly different from zero in t-tests, as reported in the Results. Nevertheless, as both isolates displayed clear evidence of substrate hydrolysis (Supplementary Figure S1), they were considered sialidase-positive in subsequent analyses. Binary logistic regression (using the packages caret [78] and pROC [79] for model evaluation) was used to evaluate the relationship between the presence of the different *nanH* genes and the probability of positive sialidase activity (as binary variable).

We investigated whether the GC-content varies between *cpn60* and *nanH1* across phylogenetic clades with a linear mixed model using gene identity, *cpn60* -clade identity and their interaction as fixed effects, and isolate identity as a random factor. We used a similar approach to analyse nucleotide composition patterns. Pairwise distances were estimated using Tamura and Nei (1993) [80] model of evolution with gamma correction as implemented in the ape package [81]. At this step, we removed two isolates from our dataset (3A6 and 3G3) because they had very short sequences due to technical issues with sequencing. Normalized *nanH1* pairwise distances were analyzed with a zero-inflated gamma model, as described above, using clade identity and the intra- vs. inter-clade factor as fixed effects, and isolate identity as a random factor.

We evaluated the link between BV-status and *cpn60* -clades, presence of *nanH* genes, and sialidase activity (as binary variable) with Fisher’s Exact Tests.

We used Kruskal–Wallis rank sum tests (implemented in the stats [69] package) to assess the effects of *cpn60* clades and BV status on MIC values for four antibiotics: augmentin, clindamycin, metronidazole, and moxifloxacin. To account for potential differences in distribution shape beyond central tendency, we confirmed the results using an Anderson–Darling k-sample test (from the kSamples [82] package). Post hoc analyzes were performed to identify pairwise differences in MIC values between *cpn60* clades using Dunn’s test for multiple comparisons (via the FSA [83] package). These analyses were further validated using pairwise Anderson–Darling tests. All post-hoc p-values were adjusted for multiple testing using the BH method.

Figures were generated using ggplot2 [84], ggpubr [85], ggmosaic [86], ggforce [87], ggExtra [88] and ggsignif [89].

## Acknowledgments

This study was funded through NRF-France Protea (UID: 116674 to R.F. and J.-A.S.P.) as well as the French embassy (Campus France) for a travel grant for V.M. to work temporarily in France. We would like to thank Yann Dumont for his help with isolation and characterization of clinical isolates in the hospital and the virostyle team at MIVEGEC for a stimulating work environment.

