## Supplementary Material for "Sialidase variability in *Gardnerella*: genetics, taxonomy, function, and clinical presentation"

January 2026

**Author Contributions:**
* These authors contributed equally to this work (co-first authors).
† These authors contributed equally to this work (co-last authors).

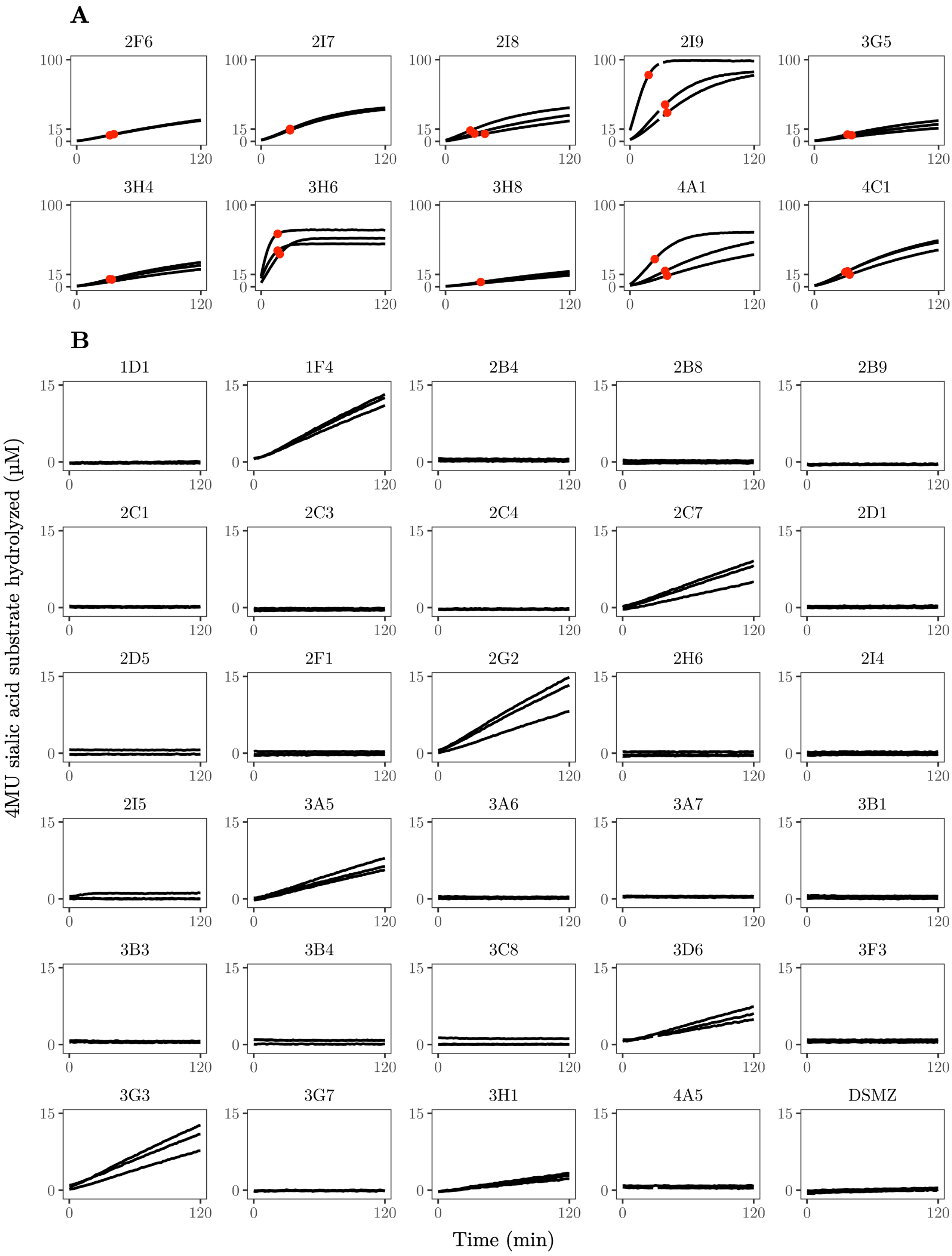


Figure S1: **Kinetics of 4MU sialic acid substrate hydrolysis for all 40 *Gardnerella* isolates.** Hydrolysis of 4MU sialic acid substrate by *Gardnerella* isolates during 120 minutes of the sialidase activity quantification assay shown as µM substrate hydrolysed at each time point. For sialidase activity estimation, the slope of the hydrolysis curve was calculated as the mean slope-value across all time points between the beginning of the assay and the beginning of the saturation, shown as a red dot (A). The absence of red dots indicate no saturation and the mean is therefore calculated over the whole assay duration (B). The three kinetics for each isolate are biological replicates (n = 3), each representing the mean of two technical replicates. Substrate hydrolysis data is based on fluorescence emitted during the assay and is calculated using 4MU standard curves as described in the methods. Note the different scales of the y-axis for panels A and B.

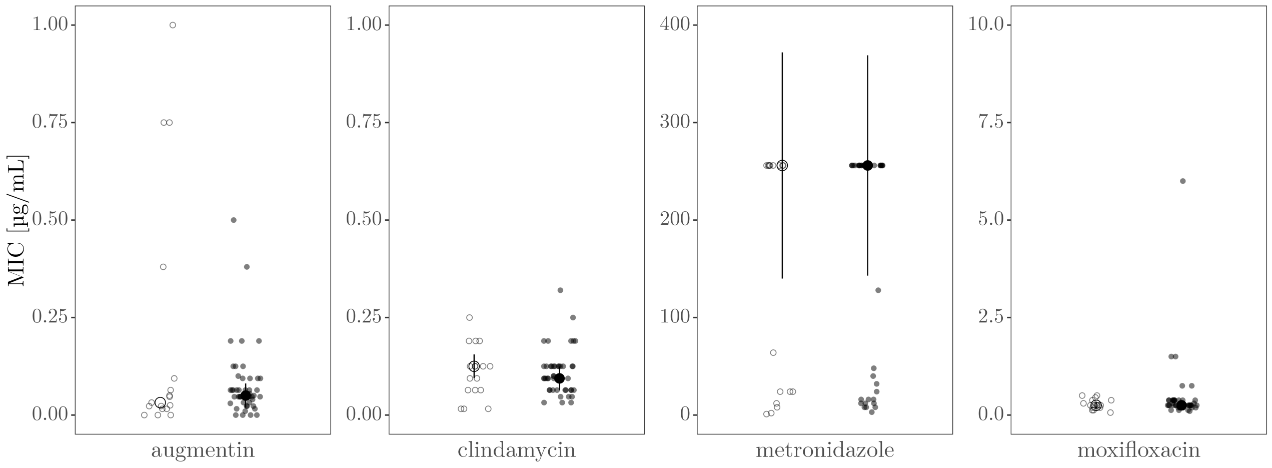


Figure S2: **Antibiotic resistance levels in clinical *Gardnerella* isolates stratified by BV status.** Individual points represent Minimum Inhibitory Concentration (MIC) values for each isolate and antibiotic. Empty points indicate BV-negative samples, while filled points indicate BV-positive samples. Larger points represent median MIC values, with error bars showing the interquartile range (IQR) around the median. Note the different y-axis scales across panels.

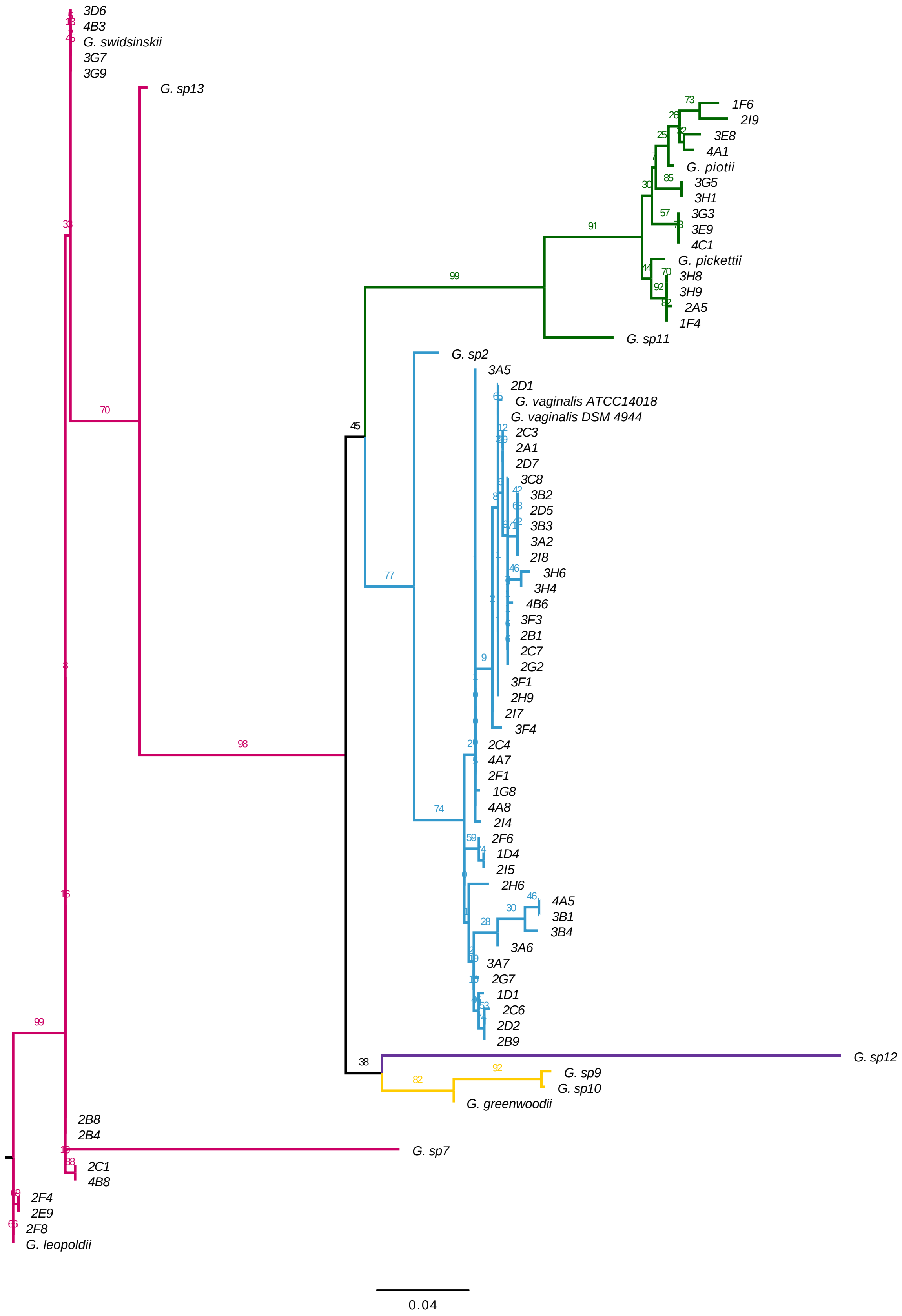


Figure S3: **Phylogenetic reconstruction of *Gardnerella* spp. isolates.** Mid-point rooted, maximum-likelihood best-known phylogenetic tree based on *cpn60* gene sequences. Branch support values are indicated and derived from 1,000 bootstrap replicates. Scale for branch length corresponds to number of nucleotide substitutions per site. Colored labels represent the five major *Gardnerella* lineages proposed for taxonomic classification.

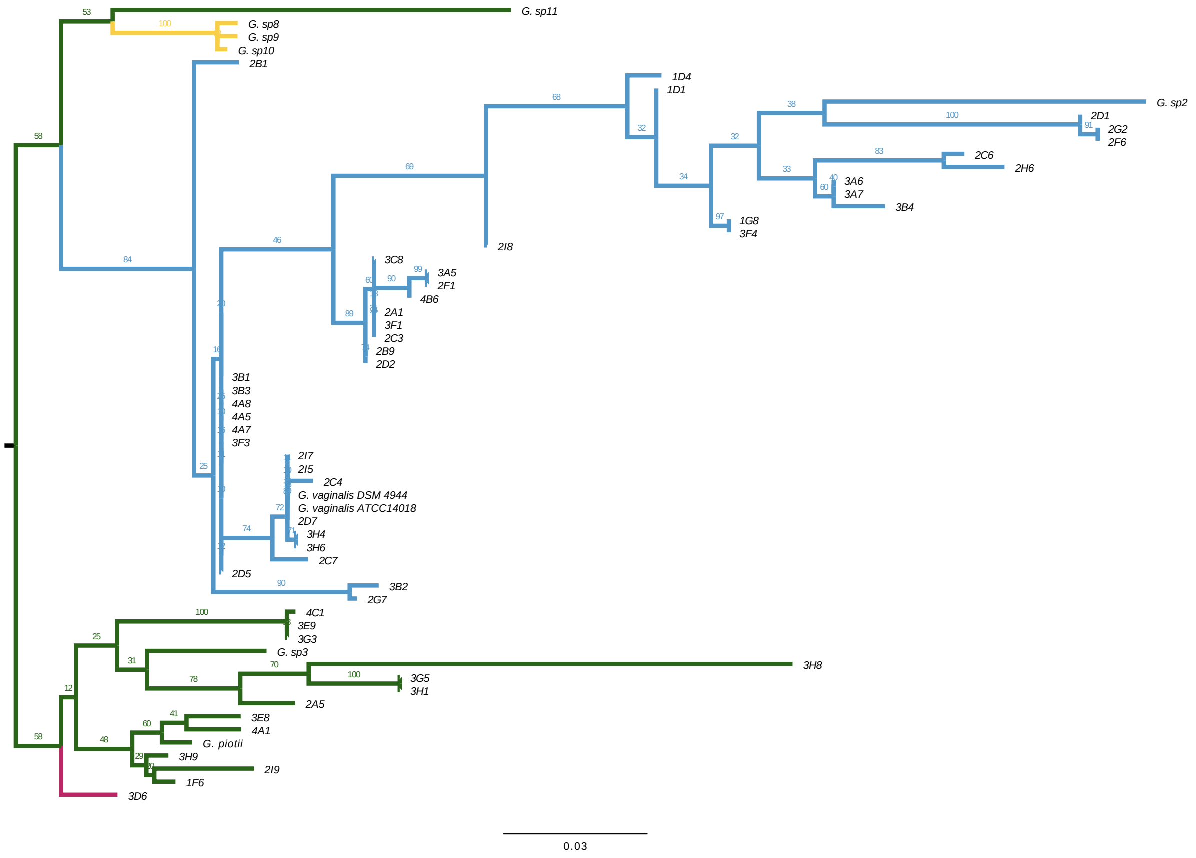


Figure S4: **Phylogenetic reconstruction of the *nanH1* gene across *Gardnerella* spp. isolates.** Midpoint rooted, maximum-likelihood best-known phylogenetic tree based on *nanH1* gene sequences. Branch support values are indicated and derived from 1,000 bootstrap replicates. Scale for branch length corresponds to number of nucleotide substitutions per site. Colored labels represent the five major *Gardnerella* lineages proposed for taxonomic classification.

Table S1: **Characteristics of *Gardnerella* isolates.** Isolates were classified as BV-positive if obtained from vaginal samples with a Spiegel's score of 3. Clade assignments based on *cpn60* and *nanH1* were inferred from maximum-likelihood phylogenetic trees constructed using the respective gene sequences. Quantitative sialidase activity is reported as the mean ± standard deviation of three biological replicates and expressed as micromoles of sialic acid hydrolyzed per minute per mg/mL of secreted protein. Sialidase gene presence was determined by PCR and antibiotic resistance is reported as minimum inhibitory concentrations (MICs, in µg/mL), measured using E-tests.

| Isolate | BV status | *cpn60* | Sialidase activity | | *nanH1* | Sialidase-gene presence | | | Antibiotic resistance | | | |
|---|---|---|---|---|---|---|---|---|---|---|---|---|
| | | clade | Qualitative | Quantitative | clade | *nanH1* | *nanH2* | *nanH3* | Metronidazole | Clindamycin | Amoxicillin | Augmentin |
| 1D1 | positive | 1 | negative | 0.006 ± 0.007 | 1 | + | - | - | 256 | 0.19 | 0.25 | 0.38 |
| 1D4 | positive | 1 | NA | NA | 1 | + | - | - | 256 | 0.25 | 1.5 | 0.125 |
| 1G8 | positive | 1 | NA | NA | 1 | + | - | + | 40 | 0.094 | 0.38 | 0.064 |
| 2A1 | negative | 1 | NA | NA | 1 | + | - | - | 24 | 0.125 | 0.25 | 0.016 |
| 2B1 | positive | 1 | NA | NA | 1 | + | - | + | 256 | 0.094 | 0.75 | 0.047 |
| 2B9 | positive | 1 | negative | 0.002 ± 0.001 | 1 | + | - | - | 128 | 0.19 | 0.25 | 0.047 |
| 2C3 | positive | 1 | negative | 0.002 ± 0.001 | 1 | + | - | - | 256 | 0.094 | 0.75 | 0.094 |
| 2C4 | positive | 1 | negative | 0.001 ± 0.002 | 1 | + | - | + | 256 | 0.064 | 0.25 | 0.047 |
| 2C6 | positive | 1 | NA | NA | 1 | + | - | - | 12 | 0.32 | 0.25 | 0.064 |
| 2C7 | positive | 1 | positive | 0.18 ± 0.049 | 1 | + | - | - | 48 | 0.064 | 6 | 0.064 |
| 2D1 | positive | 1 | negative | 0.002 ± 0.002 | 1 | + | - | - | 24 | 0.125 | 0.25 | 0.047 |
| 2D2 | positive | 1 | NA | NA | 1 | + | - | - | 256 | 0.19 | 0.38 | 0.047 |
| 2D5 | positive | 1 | negative | 0.001 ± 0.001 | 1 | + | - | - | 256 | 0.125 | 1.5 | 0.125 |
| 2D7 | negative | 1 | NA | NA | 1 | + | - | - | 256 | 0.125 | 0.5 | 0.75 |
| 2F1 | positive | 1 | negative | 0.001 ± 0.001 | 1 | + | - | + | 256 | 0.094 | 0.125 | 0.016 |
| 2F6 | positive | 1 | positive | 2.183 ± 1.147 | 1 | + | - | - | 256 | 0.125 | 0.19 | 0.19 |
| 2G2 | positive | 1 | positive | 0.433 ± 0.08 | 1 | + | - | + | 256 | 0.1 | 0.25 | 0.023 |
| 2G7 | positive | 1 | NA | NA | 1 | + | - | + | NA | NA | NA | NA |
| 2H6 | positive | 1 | negative | 0.002 ± 0.001 | 1 | + | - | - | 3 | 0.032 | 0.38 | 0.064 |
| 2H9 | negative | 1 | NA | NA | NA | - | - | - | 256 | 0.125 | 0.38 | 0 |
| 2I4 | positive | 1 | negative | 0.002 ± 0.001 | NA | - | - | - | 8 | 0.1 | 0.25 | 0.04 |
| 2I5 | positive | 1 | negative | 0.01 ± 0.018 | 1 | + | - | - | NA | 0.125 | 0.38 | 0.125 |
| 2I7 | negative | 1 | positive | 10.046 ± 9.73 | 1 | + | - | + | 64 | 0.125 | 0.5 | 0.094 |
| 2I8 | positive | 1 | positive | 2.356 ± 0.586 | 1 | + | - | + | 256 | 0.125 | 0.25 | 0.032 |
| 3A2 | negative | 1 | NA | NA | NA | + | - | - | 24 | 0.094 | 0.38 | 0.032 |
| 3A5 | negative | 1 | positive | 0.642 ± 0.177 | 1 | + | - | + | 256 | 0.19 | 0.25 | 0.05 |
| 3A6 | negative | 1 | negative | 0.001 ± 0.001 | 1 | + | + | - | 256 | 0.19 | 0.19 | 0.38 |
| 3A7 | negative | 1 | negative | 0 ± 0 | 1 | + | + | - | 2 | 0.016 | 0.125 | 0.025 |
| 3B1 | positive | 1 | negative | 0 ± 0 | 1 | + | - | - | 256 | 0.125 | 0.3 | 0.05 |
| 3B2 | positive | 1 | NA | NA | 1 | + | - | - | 256 | 0.19 | 0.38 | 0.047 |
| 3B3 | positive | 1 | negative | 0 ± 0 | 1 | + | - | - | 12 | 0.125 | 0.38 | 0.032 |
| 3B4 | positive | 1 | negative | 0 ± 0 | 1 | + | - | - | 12 | 0.094 | 0.19 | 0.047 |
| 3C8 | positive | 1 | negative | 0. 001 ± 0.001 | 1 | + | - | - | NA | NA | NA | NA |
| 3F1 | positive | 1 | NA | NA | 1 | + | - | - | 16 | 0.094 | 0.25 | 0.03 |
| 3F3 | negative | 1 | negative | 0 ± 0 | 1 | + | - | - | 24 | 0.19 | 0.3 | 0.023 |
| 3F4 | positive | 1 | NA | NA | 1 | + | - | + | 16 | 0.032 | 0.38 | 0.055 |
| 3H4 | positive | 1 | positive | 1.842 ± 0.444 | 1 | + | - | + | 256 | 0.125 | 0.38 | 0.094 |
| 3H6 | positive | 1 | positive | 13.07 ± 1.473 | 1 | + | - | + | 256 | 0.094 | 0.19 | 0.064 |
| 4A5 | negative | 1 | negative | 0.001 ± 0.002 | 1 | + | - | - | 256 | 0.064 | 0.25 | 0.064 |
| 4A7 | positive | 1 | NA | NA | 1 | + | - | - | 16 | 0.064 | 0.38 | 0.064 |
| 4A8 | NA | 1 | NA | NA | 1 | + | - | + | NA | NA | NA | NA |
| 4B6 | negative | 1 | NA | NA | 1 | + | - | + | 8 | 0.064 | 0.19 | 0.047 |
| DSMZ | positive | 1 | positive | 0.02 ± 0.003 | 1 | + | - | - | NA | NA | NA | NA |
| 1F4 | positive | 2 | positive | 0.263 ± 0.04 | NA | - | - | + | 256 | 0.032 | 0.25 | 0 |
| 1F6 | positive | 2 | NA | NA | 2 | + | - | - | 256 | 0.064 | 0.3 | 0.094 |
| 2A5 | positive | 2 | NA | NA | 2 | + | - | + | 32 | 0.064 | 0.125 | 0 |
| 2I9 | positive | 2 | positive | 8.122 ± 5.129 | 2 | + | + | + | 8 | 0.094 | 0.19 | 0 |
| 3E8 | positive | 2 | NA | NA | 2 | + | - | - | 256 | 0.125 | 0.1 | 0 |
| 3E9 | positive | 2 | NA | NA | 2 | + | - | + | 256 | 0.064 | 0.25 | 0.01 |
| 3G3 | positive | 2 | positive | 1.196 ± 0.521 | 2 | + | - | + | 256 | 0.047 | 0.38 | 0.016 |
| 3G5 | positive | 2 | positive | 1.086 ± 0.072 | 2 | + | - | + | 256 | 0.064 | 0.19 | 0.094 |
| 3H1 | positive | 2 | positive | 0.311 ± 0.123 | 2 | + | - | + | 8 | 0.064 | 0.125 | 0.064 |
| 3H8 | negative | 2 | positive | 2.188 ± 1.008 | 2 | + | - | + | 256 | 0.25 | 0.25 | 0.75 |
| 3H9 | negative | 2 | NA | NA | 2 | + | - | + | 12 | 0.094 | 0.25 | 0 |
| 4A1 | negative | 2 | positive | 6.214 ± 1.59 | 2 | + | + | - | 256 | 0.064 | 0.25 | 0 |
| 4C1 | negative | 2 | positive | 2.949 ± 0.251 | 2 | + | - | + | 1 | 0.016 | 0.064 | 0.016 |
| 2B4 | positive | 4 | negative | 0 ± 0 | NA | - | - | - | 256 | 0.047 | 0.38 | 0.023 |
| 2B8 | positive | 4 | negative | 0.001 ± 0.001 | NA | - | - | - | 256 | 0.047 | 0.25 | 0.023 |
| 2C1 | positive | 4 | negative | 0 ± 0 | NA | - | - | - | 256 | 0.125 | 0.25 | 0.047 |
| 2E9 | positive | 4 | NA | NA | NA | - | - | - | 256 | 0.125 | 0.25 | 0.064 |
| 2F4 | positive | 4 | NA | NA | NA | - | - | - | 256 | 0.19 | 0.25 | 0.5 |
| 2F8 | negative | 4 | NA | NA | NA | - | - | - | 256 | 0.125 | 0.45 | 1 |
| 3D6 | positive | 4 | positive | 0.248 ± 0.086 | 2 | + | - | + | 256 | 0.094 | 0.25 | 0.19 |
| 3G7 | negative | 4 | negative | 0.002 ± 0.001 | NA | - | - | - | 256 | 0.016 | 0.125 | 0.023 |
| 3G9 | negative | 4 | NA | NA | NA | - | - | - | NA | NA | NA | NA |
| 4B3 | positive | 4 | NA | NA | NA | - | - | - | 256 | 0.064 | 0.19 | 0.19 |
| 4B8 | positive | 4 | NA | NA | NA | - | - | - | 256 | 0.125 | 0.19 | 0.1 |

Table S2: **Metadata and relevant references of published *Gardnerella* genomespecies. .**

| ***Gardnerella* species** | **Isolate** | **cpn60 clade** | **BV status** | **Ref.** | **Sialidase activity** | **Ref.** | **Metronidazole** | **Ref.** |
|---|---|---|---|---|---|---|---|---|
| *Gardnerella vaginalis* | DSM 4944 | 1 | positive | 1 | positive | 2 | | |
| *Gardnerella vaginalis* | ATCC 14018 | 1 | positive | 1 | negative | 5 | 1, 12.5 | 6, 7 |
| *Gardnerella* sp2 | 1400E | 1 | 9 | 3 | | | 24, 32 | 6, 3 |
| *Gardnerella greenwoodii* | 00703Dmash | 3 | positive (7) | 3 & 4 | | | 256 | 3 & 6 |
| *Gardnerella* sp9 | 6119V5 | 3 | asymptomatic (5) | 3 | | | 256 | 3 & 6 |
| *Gardnerella* sp10 | 1500E | 3 | 7 | 3 | | | 256 | 3 & 6 |
| *Gardnerella piotii* | UGent 18.01 | 2 | | | positive | 5 & 6 | | |
| *Gardnerella pickettii* | 00703C2mash | 2 | positive | 4 | | | | |
| *Gardnerella* sp11 | GED7760B | 2 | | | positive | 6 | | |
| *Gardnerella leopoldii* | UGent 06.41 | 4 | | | negative | 5 | | |
| *Gardnerella swindsinskii* | GS 9838-1 | 4 | | | negative | 5 | | |
| *Gardnerella* sp7 | JCP8481A | 4 | positive | 3 | | | | |
| *Gardnerella* sp13 | KA00225 | 4 | | | | | | |
| *Gardnerella* sp12 | CMW7778B | 5 | | | | | | |

Table S3: **Association between *nanH* gene detection and BV-status.** Proportions (columns three to six) of isolates harbouring the different *nanH* genes from BV-negative and BV-positive samples. The second column shows the number of isolates in each BV-status category.

| **BV status** | **# isolates** | ***nanH1*** | ***nanH2*** | ***nanH3*** | **none** |
|---|---|---|---|---|---|
| negative | 10 | 0.90 | 0.30 | 0.40 | 0.10 |
| positive | 30 | 0.83 | 0.03 | 0.40 | 0.13 |

Table S4: **Amplicon lengths, primer sequences and PCR conditions.** PCR reagent mixes used for the PCR-reactions can be seen in Table S5.

| Gene target | Target length | Primer sequence | Primer reference | PCR mix | PCR conditions |
|---|---|---|---|---|---|
| *cpn60* | 552 bp | CGCCAGGGTTTTCCCAGTCACGAC GAIIIIGCIGGIGAYGGIACIACIAC<br>AGCGGATAACAATTTCACACAGGA YKIYKITCICCRAAICCIGGIGCYTT | H729: Jayaprakash *et al.*, 2012<br>H730: Jayaprakash *et al.*, 2012 | 2 | 94 °C for 5 min -> 40 cycles of 94 °C for 30 s, 50 °C for 60 s and 72 °C for 60 s -> 72 °C for 10 min |
| *nanH1* | 682 bp | ATGGAACGTCGTTCAACGAAG<br>GATACGCGTTTTATGTCTCTTGC | Sia1 forward: Pleckaityte *et al.*, 2012<br>Sia1 reverse: Pleckaityte *et al.*, 2012 | 1 | 94 °C for 3 min -> 40 cycles of 94 °C for 30 s, 52 °C for 60 s and 72 °C for 60 s -> 72 °C for 10 min |
| *nanH2* | 348 bp | AGGAGTGCGTATGCGTAGAAG<br>CCGCACTGCTGAGTTTCAC | G. vag nanH2 qPCR F: Robinson *et al.*, 2019<br>G. vag nanH2 qPCR R: Robinson *et al.*, 2019 | 2 | 94 °C for 3 min -> 28 cycles of 94 °C for 30 s, 52 °C for 30 s and 72 °C for 50 s -> 72 °C for 10 min |
| *nanH3* | 322 bp | CAGTTCCAATGGAAGTGTGC<br>AGCATCTGGGAATGCTCTTG | G. vag nanH3 qPCR F: Robinson *et al.*, 2019<br>G. vag nanH3 qPCR R: Robinson *et al.*, 2019 | 2 | 94 °C for 3 min -> 28 cycles of 94 °C for 30 s, 52 °C for 30 s and 72 °C for 50 s -> 72 °C for 10 min |

Table S5: **PCR reagent mixes.** We used 20 µL of DNA template for Mix 1 and 10 µL of template for mix 2. Mixes 1 and 2 were filled up to the final reaction volume (50 µL and 25 µL, respectively) with Milli-Q water. The final primer concentration in PCR Mix 2 was 400 nM for *cpn60* amplifications and 500 nM for all PCRs of *nanH*-genes.

| **Mix 1** | | **Mix 2** | |
|---|---|---|---|
| Reagent | Final concentration | Reagent | Final concentration |
| 10X PCR reaction buffer | 1X | 10X PCR reaction buffer | 1X |
| 50 mM Magnesium chloride ($MgCl_2$) | 1 mM (total of 2.5 mM with $MgCl_2$ from reaction buffer) | 50 mM Magnesium chloride ($MgCl_2$) | 1 mM (total of 2.5 mM with $MgCl_2$ from reaction buffer) |
| 10 mM PCR grade nucleotide mix | 250 µM of each dNTP | 10 mM PCR grade nucleotide mix | 200 µM of each dNTP |
| Taq DNA polymerase (Sigma-Aldrich, TAQN-RO) | 2.5 units | Taq DNA polymerase (Sigma-Aldrich, TAQN-RO) | 1.5 units |
| | | DMSO | 5% (v/v) |
| Primer mix | 800 nM | Primer mix | 400 or 500 nM |
| **Final volume** | 50 µl | **Final volume** | 25 µl |

Table S6: **NCBI accession numbers and cpnID of *Gardnerella* spp. reference sequences.** All *Gardnerella* spp. strains that were used as reference sequences for the 13 genome species described by Vaneechoutte *et al.* (2019) with their NCBI accession numbers and their corresponding ID in the cpn database.

| ***Gardnerella* species** | **NCBI accession** | **cpnID** |
|---|---|---|
| *Gardnerella leopoldii* UGent 06.41 | GCA_003293675.1 | b32837 |
| *Gardnerella piotii* UGent 18.01 | QJUV00000000 | b32843 |
| *Gardnerella swidsinskii* GS 9838-1 | QJVB00000000 | b32838 |
| *Gardnerella vaginalis* ATCC 14018 | SJWZ00000000 | b291 |
| *Gardnerella pickettii* 00703C2mash | ADEU00000000 | b21783 |
| *Gardnerella greenwoodii* 00703Dmash | ADEV00000000 | b21782 |
| *Gardnerella*-1400E-sp2 | ADER00000000 | b21786 |
| *Gardnerella*-JCP8481A-sp7 | ATJG00000000 | b26734 |
| *Gardnerella*-6119V5-sp9 | ADEW00000000 | b21781 |
| *Gardnerella*-1500E-sp10 | ADES00000000 | b21785 |
| *Gardnerella*-GED7760B-sp11 | LRQA00000000 | b28176 |
| *Gardnerella*-CMW7778B-sp12 | LSRC00000000 | b28178 |
| *Gardnerella*-KA00225-sp13 | MNLH00000000 | b32839 |

Table S7: **Optical density measurements of low density samples.** For some replicates (indicated in the second column of the table), sample densities as estimated by optical density ($OD_{600}$) measurements were too low to be adjusted to $OD_{600}$ of 0.8 after growth on SB agar. In all cases, included those here reported, sialidase activity quantification was normalized by total protein concentration.

| **Isolate** | **Replicate** | $\mathbf{OD_{600}}$ |
|---|---|---|
| 2I9 | 1 | 0.724 |
| 3A1 | 2 | 0.681 |
| 3A1 | 3 | 0.65 |
| 3D6 | 3 | 0.705 |
| 1I4 | 1 | 0.423 |
| 1I4 | 2 | 0.399 |
| 1I4 | 3 | 0.449 |
| 4A5 | 3 | 0.749 |
| 2B4 | 1 | 0.717 |
| 2B4 | 2 | 0.761 |
| 3A7 | 1 | 0.734 |
| 3B3 | 3 | 0.695 |
| 3B4 | 2 | 0.684 |
| 3C8 | 2 | 0.771 |
| 3F3 | 1 | 0.768 |
| 3F3 | 2 | 0.671 |
| 3G5 | 2 | 0.722 |
| 3G5 | 3 | 0.669 |
| 3H6 | 1 | 0.669 |
| 3H6 | 2 | 0.62 |
| 4A1 | 1 | 0.666 |
| 4A1 | 3 | 0.686 |
| 4B9 | 1 | 0.47 |
| 4B9 | 2 | 0.568 |
| 4B9 | 3 | 0.621 |

Table S8: **Antibiotic resistance test results.** Summary of statistical analyses comparing median MIC values across different *Gardnerella* clades and BV statuses. Differences were assessed using Kruskal–Wallis tests (with Dunn's post hoc comparisons) and Anderson–Darling k-sample tests, shown side by side for comparison. Significance levels: * = $p < 0.05$, ** = $p < 0.01$, *** = $p < 0.001$, ns = Not significant ($p \geq 0.05$).

Median and interquartile range (IQR) of MIC values (µg/mL) per antibiotic type and *cpn60* clade

| Antibiotic type | Clade | Median | IQR |
|---|---|---|---|
| Augmentin | 1 | 0.05 | 0.02 |
| Augmentin | 2 | 0.01 | 0.03 |
| Augmentin | 4 | 0.08 | 0.08 |
| Clindamycin | 1 | 0.13 | 0.02 |
| Clindamycin | 2 | 0.06 | 0.02 |
| Clindamycin | 4 | 0.11 | 0.04 |
| Metronidazole | 1 | 256 | 119 |
| Metronidazole | 2 | 256 | 122 |
| Metronidazole | 4 | 256 | 0 |
| Moxifloxacin | 1 | 0.30 | 0.07 |
| Moxifloxacin | 2 | 0.25 | 0.06 |
| Moxifloxacin | 4 | 0.25 | 0.02 |

Kruskal-Wallis test; global differences

| Antibiotic type | Predictor | Chi.squared | Df | P.value | Significance |
|---|---|---|---|---|---|
| Augmentin | BV status | 0.498 | 1 | 0.481 | ns |
| Augmentin | Clade | 8.742 | 2 | 0.013 | * |
| Clindamycin | BV status | 0.044 | 1 | 0.835 | ns |
| Clindamycin | Clade | 7.703 | 2 | 0.021 | * |
| Metronidazole | BV status | 1.033 | 1 | 0.309 | ns |
| Metronidazole | Clade | 6.646 | 2 | 0.036 | * |
| Moxifloxacin | BV status | 0.262 | 1 | 0.609 | ns |
| Moxifloxacin | Clade | 9.190 | 2 | 0.010 | * |

Anderson-Darling k-sample test; global differences

| Antibiotic type | Predictor | T.AD | P.value | Significance |
|---|---|---|---|---|
| Augmentin | BV status | 0.921 | 0.135 | ns |
| Augmentin | Clade | 7.278 | < 0.001 | *** |
| Clindamycin | BV status | -0.080 | 0.391 | ns |
| Clindamycin | Clade | 3.214 | 0.013 | * |
| Metronidazole | BV status | -0.008 | 0.360 | ns |
| Metronidazole | Clade | 3.979 | 0.006 | ** |
| Moxifloxacin | BV status | -0.805 | 0.860 | ns |
| Moxifloxacin | Clade | 4.274 | 0.004 | ** |

Dunn's post-hoc test; pairwise comparisons between clades

| Antibiotic type | Clade comparison | Z | P.unadj | P.adj | Significance |
|---|---|---|---|---|---|
| Augmentin | 1 - 2 | 2.537 | 0.011 | 0.017 | * |
| Augmentin | 1 - 4 | -0.907 | 0.365 | 0.365 | ns |
| Augmentin | 2 - 4 | -2.695 | 0.007 | 0.021 | * |
| Clindamycin | 1 - 2 | 2.725 | 0.006 | 0.019 | * |
| Clindamycin | 1 - 4 | 1.128 | 0.259 | 0.389 | ns |
| Clindamycin | 2 - 4 | -1.124 | 0.261 | 0.261 | ns |
| Metronidazole | 1 - 2 | -0.142 | 0.887 | 0.887 | ns |
| Metronidazole | 1 - 4 | -2.537 | 0.011 | 0.033 | * |
| Metronidazole | 2 - 4 | -2.036 | 0.042 | 0.063 | ns |
| Moxifloxacin | 1 - 2 | 2.896 | 0.004 | 0.011 | * |
| Moxifloxacin | 1 - 4 | 1.528 | 0.127 | 0.190 | ns |
| Moxifloxacin | 2 - 4 | -0.917 | 0.359 | 0.359 | ns |

Pairwise Anderson-Darling tests; pairwise comparisons between clades

| Antibiotic type | Clade comparison | P.unadj | P.adj | Significance |
|---|---|---|---|---|
| Augmentin | 1 - 2 | 0.000 | 0.001 | ** |
| Augmentin | 1 - 4 | 0.187 | 0.187 | ns |
| Augmentin | 2 - 4 | 0.003 | 0.004 | ** |
| Clindamycin | 1 - 2 | 0.002 | 0.005 | ** |
| Clindamycin | 1 - 4 | 0.396 | 0.411 | ns |
| Clindamycin | 2 - 4 | 0.411 | 0.411 | ns |
| Metronidazole | 1 - 2 | 0.821 | 0.821 | ns |
| Metronidazole | 1 - 4 | 0.001 | 0.003 | ** |
| Metronidazole | 2 - 4 | 0.005 | 0.008 | ** |
| Moxifloxacin | 1 - 2 | 0.001 | 0.004 | ** |
| Moxifloxacin | 1 - 4 | 0.138 | 0.207 | ns |
| Moxifloxacin | 2 - 4 | 0.423 | 0.423 | ns |

Table S9: **Published datasets on sialidase activity of *Gardnerella* spp. isolates.** Summary information of published datasets that provide sialidase activity data of *Gardnerella* spp. clinical isolates including information on taxonomy, BV status or the *nanH* gene repertoire (*nanH1-3*).

| Study | Taxonomy | BV status | Type of sialiadase data | Qualitative sialidase activity | *nanH1* | *nanH2* | *nanH3* |
|---|---|---|---|---|---|---|---|
| Briselden *et al.* 1992 | n = 41<br>na | neg: 46.3%<br>pos: 53.7% | quantitative<br>unit = µmoles 4MU per mg protein per min | BV- : 3.5 ± 3.6 U<br>BV+ : 12.0 ± 8.1 U | na | na | na |
| Lopes dos Santos Santiago *et al.* 2011 | n = 33<br>ARDRA1: 21.21%<br>ARDRA2: 42.42%<br>ARDRA3: 36.36% | neg: 0%<br>int: 21.21%<br>pos: 72.72% | qualitative<br>filter spot test | ARDRA1: 100% (n = 7)<br>ARDRA2: 0% (n = 14)<br>ARDRA3: 100% (n = 12)<br>BV- : na<br>BVint : 85.71% (n = 7)<br>BV+ : 70.83% (n = 24) | ARDRA1: 100% (n = 7)<br>ARDRA2: 0% (n = 14)<br>ARDRA3: 100% (n = 12)<br>BV- : na<br>BVint : 85.71% (n = 7)<br>BV+ : 70.83% (n = 24) | na | na |
| Pleckaityte *et al.* 2012 | n = 17<br>ARDRA1: 64.7%<br>ARDRA2A: 5.9%<br>ARDRA2AB: 11.76%<br>ARDRA2B: 5.9%<br>ARDRA2C: 11.76% | na | qualitative<br>filter spot test | ARDRA1: 36.36% (n = 11)<br>ARDRA2A: 100% (n = 1)<br>ARDRA2AB: 100% (n = 2)<br>ARDRA2B: 100% (n = 1)<br>ARDRA2C: 100% (n = 2) | ARDRA1: 100% (n = 11)<br>ARDRA2A: 100% (n = 1)<br>ARDRA2AB: 100% (n = 2)<br>ARDRA2B: 100% (n = 1)<br>ARDRA2C: 100% (n = 2) | na | na |
| Lewis *et al.* 2013 | n = 15<br>na | neg: 13.33%<br>int: 6.67%<br>pos: 80% | quantitative<br>unit = sialidase activity normalized by OD | BV- : 100% (n = 2)<br>BVint : 100% (n = 1)<br>BV+ : 66.67% (n = 12) | na | na | na |
| Castro *et al.* 2015 | n = 6<br>na | neg: 50%<br>pos: 50% | quantitative<br>unit = normalized expression in relation to 16S rRNA | BV- : 100% (n = 3)<br>BV+ : 100% (n = 3) | BV- : 85.71% (n = 7)<br>BV+ : 71.43% (n = 7) | na | na |
| Schellenberg *et al*. 2016<br>(*nanH3* data from Kurukulasuriya *et al.* 2021) | n = 112<br>1: 31.25%<br>2: 29.46%<br>3: 7.14%<br>4: 32.14% | na | quantitative<br>unit = ΔRFU/min normalized by OD | 1: 8.3% (n = 35)<br>2: 100% (n = 33)<br>3: 0% (n = 8)<br>4: 0% (n = 36) | 1: 100% (n = 35)<br>2: 100% (n = 33)<br>3: 100% (n = 8)<br>4: 2.8% (n = 36) | na | 1: 8.57% (n = 35)<br>2: 96.97% (n = 33)<br>3: 0% (n = 8)<br>4: 0% (n = 36) |
| Vaneechoutte *et al.* 2019 | n = 10<br>1: 40%<br>2: 20%<br>3: 0%<br>4: 40% | na | qualitative<br>filter spot test | 1: 25% (n = 4)<br>2: 100% (n = 2)<br>3: na<br>4: 0% (n = 4) | na | na | na |
| Robinson *et al.* 2019 | n = 34<br>na | na | quantitative<br>unit = ΔRFU/min | not specified by taxonomy or BV status<br>41.18% | 91.18% | 14.71% | 41.18% |
| Bulavaitė *et al.* 2021<br>(quantitative sialidase activity data from Janulaitiene *et al.* 2018) | n = 34<br>1: 44.12%<br>2: 29.41%<br>3: 0%<br>4: 26.47% | na | quantitative<br>unit = ΔRFU/min normalized by OD | 1: 60% (n = 15)<br>2: 100% (n = 10)<br>3: na<br>4: 0% (n = 9) | 1: 100% (n = 15)<br>2: 100% (n = 10)<br>3: na<br>4: 0% (n = 9) | 1: 0% (n = 15)<br>2: 40% (n = 10)<br>3: na<br>4: 0% (n = 9) | 1: 26.67% (n = 15)<br>2: 90% (n = 10)<br>3: na<br>4: 0% (n = 9) |
| Shipitsyna *et al.* 2022 | n = 19<br>1: 84.21%<br>2: 0%<br>3: 0%<br>4: 15.79% | na | semi-quantitative<br>four categories of fluorescence data | 1: 37.5% (n = 16)<br>2: na<br>3: na<br>4: 0% (n = 3) | 1: 100% (n = 16)<br>2: na<br>3: na<br>4: 0% (n = 3) | 1: 0% (n = 16)<br>2: na<br>3: na<br>4: 0% (n = 3) | 1: 43.75% (n = 16)<br>2: na<br>3: na<br>4: 0% (n = 3) |
| Sousa *et al.* 2023 | n = 2<br>1: 0%<br>2: 50%<br>3: 50%<br>4: 0% | na | qualitative<br>filter spot test | 1: na<br>2: 100% (n = 1)<br>3: 0% (n = 1)<br>4: na | 1: na<br>2: 100% (n = 1)<br>3: 100% (n = 1)<br>4: na | 1: na<br>2: 0% (n = 1)<br>3: 0% (n = 1)<br>4: na | 1: na<br>2: 100% (n = 1)<br>3: 0% (n = 1)<br>4: na |
| This study | n = 66<br>1: 63.64%<br>2: 19.7%<br>3: 0%<br>4: 16.66%<br>5: 0% | neg: 27.27%<br>pos: 71.21% | quantitative<br>unit = µmoles 4MU per mg protein per min | 1: 30.77% (n = 26)<br>2: 100% (n = 8)<br>3: na<br>4: 20% (n = 5)<br>5: na<br>BV- : 50% (n = 10)<br>BV+ : 41.38% (n = 29) | 1: 95.24% (n = 42)<br>2: 92.31% (n = 13)<br>3: na<br>4: 9.09% (n = 11)<br>5: na<br>BV- : 77.78% (n = 18)<br>BV+ : 80.85% (n = 47) | 1: 4.76% (n = 42)<br>2: 15.38% (n = 13)<br>3: na<br>4: 0% (n = 11)<br>5: na<br>BV- : 0.17% (n = 18)<br>BV+ : 0.02% (n = 47) | 1: 33.33% (n = 42)<br>2: 76.92% (n = 13)<br>3: na<br>4: 9.09% (n = 11)<br>5: na<br>BV- : 0.33% (n = 18)<br>BV+ : 38.30% (n = 47) |
